# *De novo* peptide databases enable protein-based stable isotope probing of microbial communities with up to species-level resolution

**DOI:** 10.1101/2024.11.25.625156

**Authors:** Simon Klaes, Christian White, Lisa Alvarez-Cohen, Lorenz Adrian, Chang Ding

## Abstract

**Background:** Protein-based stable isotope probing (Protein-SIP) is a powerful approach that can directly link individual taxa to activity and substrate assimilation, elucidating metabolic pathways and trophic relationships within microbial communities. In Protein-SIP, peptides and corresponding taxa are identified by database matching, making database quality crucial for accurate analyses. For samples with unknown community composition, Protein-SIP typically employs either unrestricted reference databases or metagenome-derived databases. While (meta)genome-derived databases represent the gold standard, they may be incomplete and are typically resource-intensive to generate. In contrast, unrestricted reference databases can inflate the search space and require complex post-processing.

**Results:** Here, we explore the feasibility of using *de novo* peptide sequencing to construct peptide databases directly from mass spectrometry raw data. We then use the mass spectrometric data from labeled cultures to quantify isotope incorporation into specific peptides. We benchmark our approach against the canonical approach in which a sample-matching (meta)genome-derived protein sequence database is used on three different datasets: 1) a proteome analysis from a defined microbial community containing ^13^C-labeled *E. coli* cells, 2) time-course data of an anammox-dominated continuous reactor after feeding with ^13^C-labeled bicarbonate, and 3) a model of the human distal gut simulating a high-protein and high-fiber diet cultivated in either ^2^H2O or H2^18^O. Our results show that *de novo* peptide databases are applicable to different isotopes, detecting similar amounts of labeled peptides compared to sample-matching (meta)genome-derived databases, and also identify labeled peptides missed by this canonical approach. Furthermore, we show that peptide-centric Protein-SIP allows up to species-specific resolution and enables the assessment of activity related to individual biological processes. Finally, we provide access to our modular Python pipeline to assist the construction of *de novo* peptide databases and subsequent peptide-centric Protein-SIP data analysis (https://git.ufz.de/meb/denovo-sip).

**Conclusions:** *De novo* peptide databases enable Protein-SIP of microbial communities without prior knowledge of the composition and can be used complementarily to (meta)genome-derived databases or as a standalone alternative in exploratory or resource-limited settings.

## Introduction

Microbial communities are ubiquitous and crucial for the functioning and health of myriad ecosystems. Thus, understanding microbial communities is essential for elucidating fundamental ecological processes and advancing knowledge of human health and the environment [1–3]. Microbial communities are frequently studied via omics approaches [4], which can provide insights into their state and dynamics [5]. However, the complexity of microbial communities often hinders attempts to link ecosystem functions to individual taxa. To establish direct links between substrates, metabolites, functions, and taxa, omics approaches can be combined with stable isotope probing (SIP) [6].

In SIP experiments, microbial communities are exposed to a substrate enriched in a stable isotope. Upon assimilation, microorganisms can incorporate the stable isotopes into biomolecules, such as DNA [7, 8], RNA [8, 9], phospholipid-derived fatty acids [10, 11], and proteins [12, 13]. Analyzing the relative amount of labeled biomolecules and their degree of isotope incorporation can establish a direct link of individual taxa to general (for ^2^H or ^18^O labeled water) or substrate-specific (for ^13^C, ^15^N, or ^34^S labeled substrate) activity [8–19]. By establishing this link, SIP can identify key microorganisms [19, 20], elucidate metabolic pathways [21, 22], and unravel complex food webs in microbial communities [8, 23]. The choice of the investigated biomolecule affects the sensitivity and taxonomic resolution of SIP analyses (as reviewed in [24]). DNA and RNA-based SIP provide high taxonomic resolution but rely on the separation of heavy and light fractions via centrifugation, leading to inaccuracy in isotope quantification and low sensitivity, typically requiring isotope enrichments of >10 atom% for RNA and >20 atom% for DNA [25, 26]. Fatty acid and protein-based SIP (Protein-SIP) employ mass spectrometry to accurately quantify isotope incorporation down to 0.01 atom% isotope enrichment [27]. While fatty acids only provide limited taxonomic information, proteins can characterize communities often at the species level [28], and, in some cases, even at the strain level [29]. Thus, Protein-SIP offers the advantage of accurately quantifying isotope incorporation while preserving high taxonomic resolution.

Protein-SIP is based on bottom-up metaproteomics. In brief, proteins are extracted from the sample and digested into peptides that are then analyzed by high-resolution tandem mass spectrometry (MS/MS). Obtained spectra are mapped back to peptides and relative isotope abundances (RIA) are calculated from shifts in precursor ion peaks [30–32]. In principle, spectra can be mapped to peptides by database or spectral library searches, or sequenced by *de novo* peptide sequencing. To date, only database searches are used for Protein-SIP. In database searches, acquired mass spectra are compared to theoretical spectra of peptides in a protein or peptide sequence database to obtain peptide-spectrum matches (PSMs) that are then validated by a target-decoy approach. While database-dependent approaches are widely accepted for proteomics and Protein-SIP, they are heavily reliant on the quality of the database leading to several constraints. One key constraint is that database creation ideally requires prior knowledge of the community composition. For defined cultures, such as pure strains, sample-matching databases are readily available in public repositories. However, for samples of unknown composition, *e.g.*, environmental samples, databases usually must be derived from metagenome sequencing. Metagenome-derived protein sequence databases are considered the gold standard as they enable high peptide identification rates and offer high taxonomic resolution [33]. However, generating metagenome-derived protein sequence databases is resource-intensive, time-consuming, and requires specialized expertise. A common challenge in this process is that a substantial portion of metagenomic reads typically remains unassembled, assembled contigs often cannot be confidently binned into genomes, and the resulting metagenome-assembled genomes are frequently incomplete [34–38]. Despite above-mentioned challenges, metagenome-derived protein sequence databases typically represent the best choice for metaproteomic analyses of samples with unknown composition [33]. In cases where metagenomes are not available, unrestricted reference databases such as NCBI non-redundant proteins (nr) or Uniprot can be used [12, 39–42]. However, unrestricted reference databases substantially increase computational time, require advanced filtering strategies, and often lead to lower taxonomic resolution [43–46]. Subsets of unrestricted reference databases focusing on taxonomic marker proteins were successfully applied to Protein-SIP, reducing computational time, but also limiting the taxonomic resolution [47]. Both, metagenome-derived databases and unrestricted reference databases contain sequences of proteins that are not expressed in the sample, inflating database size. Inflated databases can negatively impact peptide identification during target-decoy validation strategies [44, 48]. Furthermore, database-dependent search approaches are inherently biased towards the database content making it impossible to detect proteins missing in the database [45].

*De novo* peptide sequencing (hereafter: *de novo* sequencing) addresses the above-mentioned limitations of database-dependent searches by directly inferring peptides from MS/MS spectra without using references. Thereby, *de novo* peptide sequencing does not provide a link of peptides to proteins like database-dependent approaches, making it a peptide-centric approach, in which taxonomic and functional information of peptides can be inferred by text matching (Unipept [49]) or sequence similarity (BLASTP [50]). To date, *de novo* sequencing has been used to characterize microbial community compositions at the family level in complex communities [51] and at the species level in pure cultures or simple communities [52]. Until recently, *de novo* sequencing identified far fewer peptides than database-dependent approaches [53] and faced inherent challenges in controlling false discovery rates (FDRs). However, current advances in artificial intelligence and the increased amount of publicly available training data have massively improved *de novo* sequencing algorithms, challenging database-dependent approaches [54, 55]. Additionally, hybrid methods that subject *de novo* identified peptides to subsequent database searches may control FDRs [56, 57].

Despite the advances in *de novo* sequencing, it remains unclear whether *de novo* algorithms can reliably identify peptides in partially labeled samples, where isotope peaks may cause signal interference. Moreover, *de novo* sequencing alone cannot identify labeled peptides or quantify isotope incorporation, necessitating their integration with SIP tools. To address this knowledge gap, we propose the creation of sample-matching databases derived from *de novo* sequencing of unlabeled peptides, enabling Protein-SIP analysis without prior knowledge of community compositions (Figure 1).

**Figure 1:**
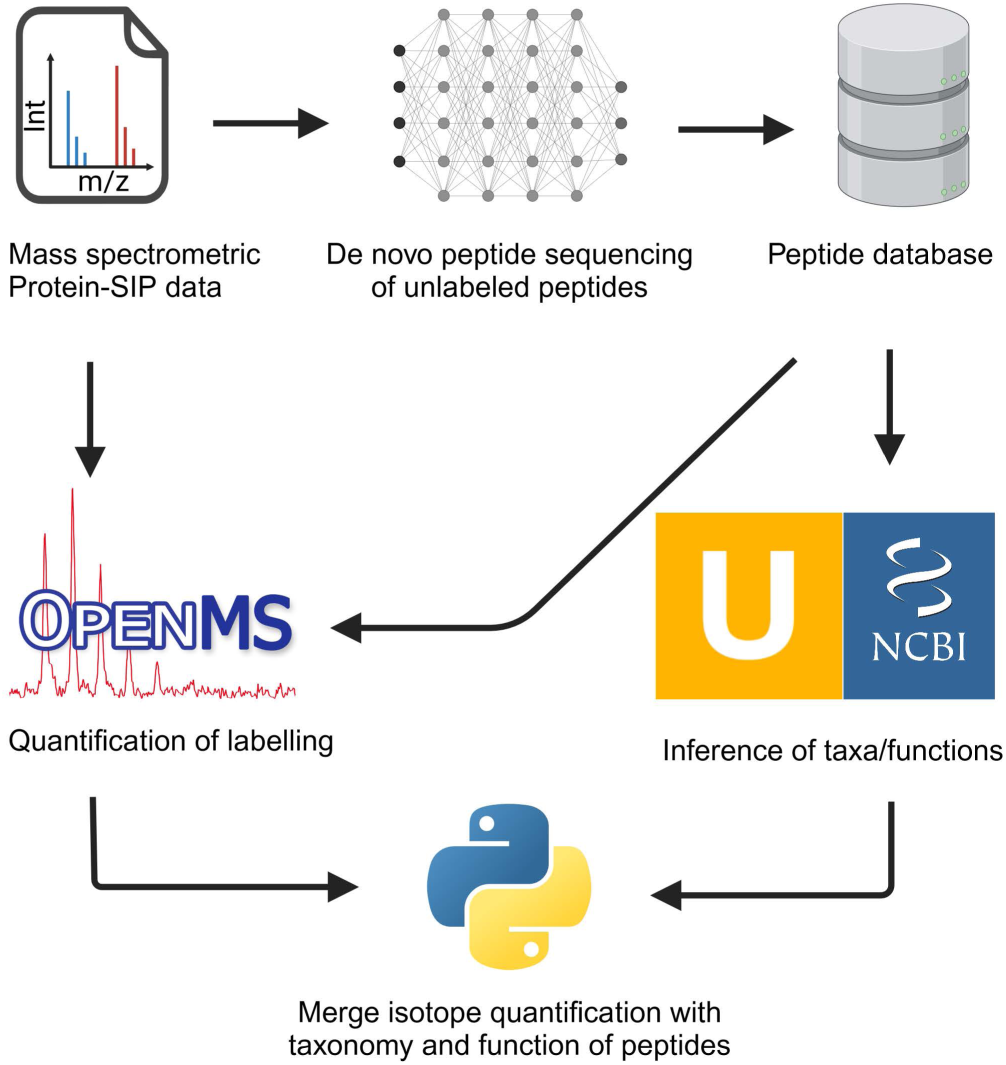
Proposed workflow for protein-based stable isotope probing (Protein-SIP) using *de novo* peptide databases. Here, we investigate whether *de novo* algorithms can accurately identify peptides in partially labeled samples and assess whether these identified peptides can be used to assemble sample-matching databases for Protein-SIP analysis using ^2^H, ^13^C, and ^18^O labeled data. Additionally, we benchmark peptide databases created with different state-of-the-art *de novo* sequencing tools with each other and with genome-derived databases.

## Results

### *De novo* sequencing of peptides in partially labeled samples

To investigate (1) whether *de novo* algorithms can identify peptides in partially labeled samples, and (2) how the RIA affects *de novo* peptide sequencing, we analyzed previously published proteomic data (PRIDE accession number: PXD04141) of standard *E. coli* cultures cultivated with a defined ^13^C RIA of 1.07–99% [32]. Specifically, we performed *de novo* peptide sequencing using the state-of-the-art algorithms PepNet and Casanovo and quantified the number of identified peptides classified as either matching or not matching the *E. coli* K12 reference proteome or common contaminants across the reported quality scores. For comparison, we also performed conventional database search using MS-GF+ with a *E. coli* K12 reference protein database from UniProt (PRIDE accession number: PXD024285) supplemented with common contaminant sequences (total database size: 4,694 protein sequences, Figure 2).

**Figure 2:**
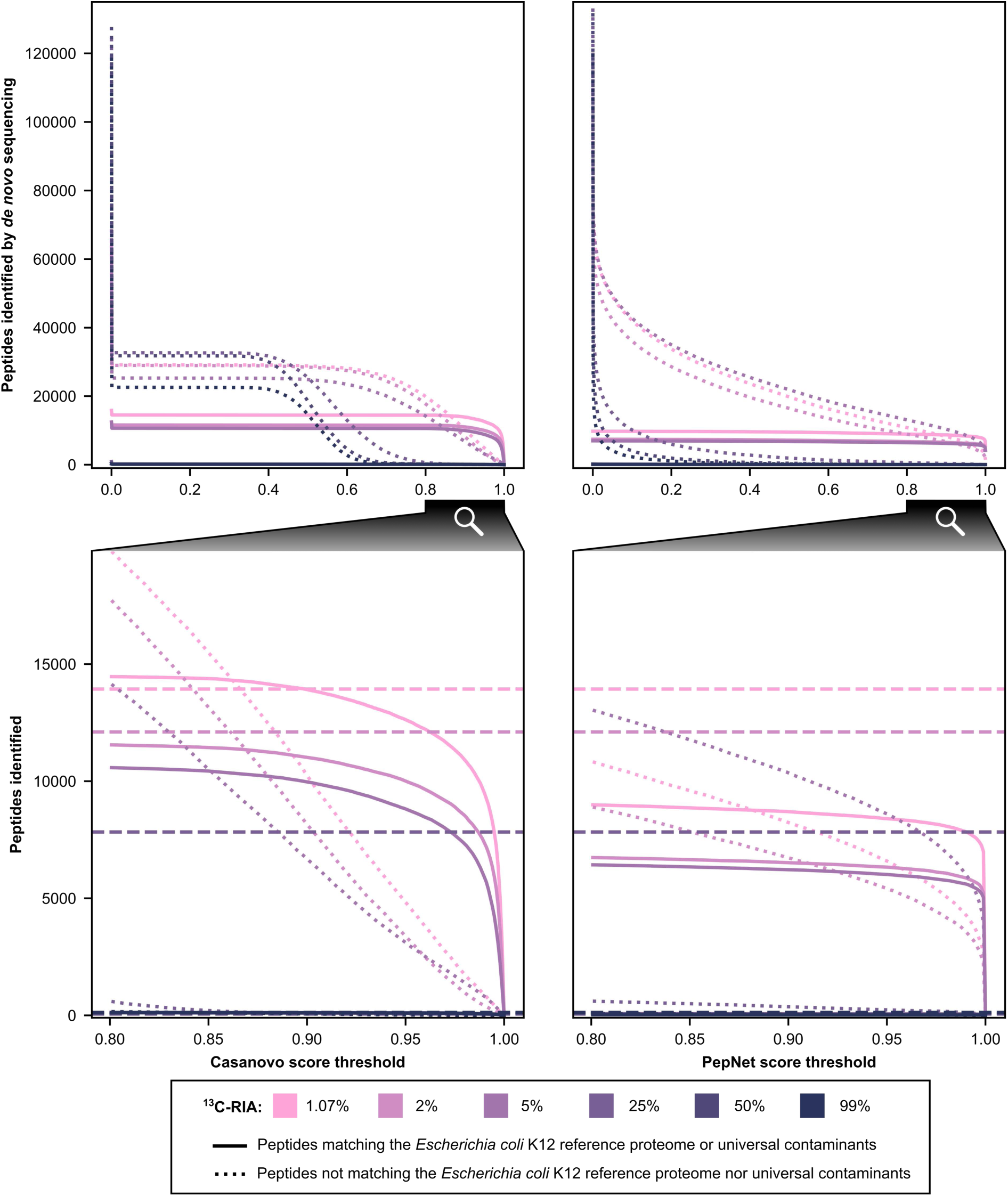
Comparison of the non-redundant number and quality score of peptides identified by *de novo* sequencing in standard *Escherichia coli* K12 cultures across biological triplicates cultivated with a defined ^13^C relative isotope abundance (RIA) of 1.07–99%. *De novo* peptide sequencing was performed using Casanovo and PepNet. Negative quality scores are plotted as zero. Solid lines represent identified peptides matching the proteome of *E. coli* K12 or universal contaminants. Dotted lines indicate identified peptides not matching the proteome of *E. coli* K12 nor universal contaminants. Dashed lines are only plotted in the bottom graphs and represent the number of identifications using the genome-derived reference protein database with MS-GF+ at 1% PSM-level FDR.

Without filtering for any score threshold, Casanovo and PepNet successfully identified 6,956–15,824 unique *E. coli* K12 and universal contaminant peptides in samples containing a ^13^C RIA of 1.07–5% ^13^C. An increase in the ^13^C RIA resulted in a reduced number of peptide identifications. In samples with a ^13^C RIA of 25–99%, both *de novo* algorithms as well as conventional database search failed to detect a relevant number of *E. coli* K12 peptides. However, this limitation is not critical, as SIP tools usually require an unlabeled reference for detecting peptides containing more than 10% ^13^C [27]. These unlabeled references can then be used for *de novo* sequencing to create the peptide database.

Casanovo outperformed PepNet by identifying more *E. coli* K12 peptides and fewer non-*E. coli* K12 peptides. Moreover, applying higher quality score thresholds led to a pronounced decrease in non-*E. coli* K12 peptide identifications but only a marginal decrease in *E. coli* K12 peptide identifications. This indicates effective separation of true and false-positive identifications by Casanovo at high quality scores. Compared to MS-GF+ database search, Casanovo identified 103.1–157.7% as many unique *E. coli* K12 and universal contaminant peptides. However, Casanovo also reported approximately ten times more peptides not attributable to *E. coli* K12 or known contaminants than those that are attributable to *E. coli* K12 or known contaminants, indicating the need for post-processing to control the FDR.

### Benchmarking *de novo* assembled databases with a defined microbial community

To test (1) whether identified peptides can be an adequate database for identifying labeled peptides and (2) how the FDR can be controlled, we analyzed previously published proteomic data (PRIDE accession number: PXD024174) from a defined microbial community containing unlabeled and ^13^C-labeled *E. coli* cells in a 1:1 ratio [27].The community consisted of 21 bacterial species belonging to 15 families, 1 eukaryote, 1 archaeon and 5 bacterial viruses (Supplementary Table 1 in [28]). The labeled *E. coli* cells were cultivated with 10% fully labeled (100% ^13^C) and 90% unlabeled (1.1% ^13^C) glucose. We performed *de novo* peptide sequencing with PepNet and Casanovo. Subsequently, peptides were filtered applying different defined score thresholds to assemble databases of 14,700 to 875,366 peptides (Supplementary Table S1) which were then used to identify unlabeled peptides via MS-GF+ and labeled peptides via MetaProSIP (Figure 3A and Supplementary Figure S1) The metagenome-derived protein sequence database used for comparison was assembled by Kleiner et al. [28], incorporating all predicted proteins from reference genomes of the mock community members and universal contaminants, totaling 123,365 protein sequences.

**Figure 3:**
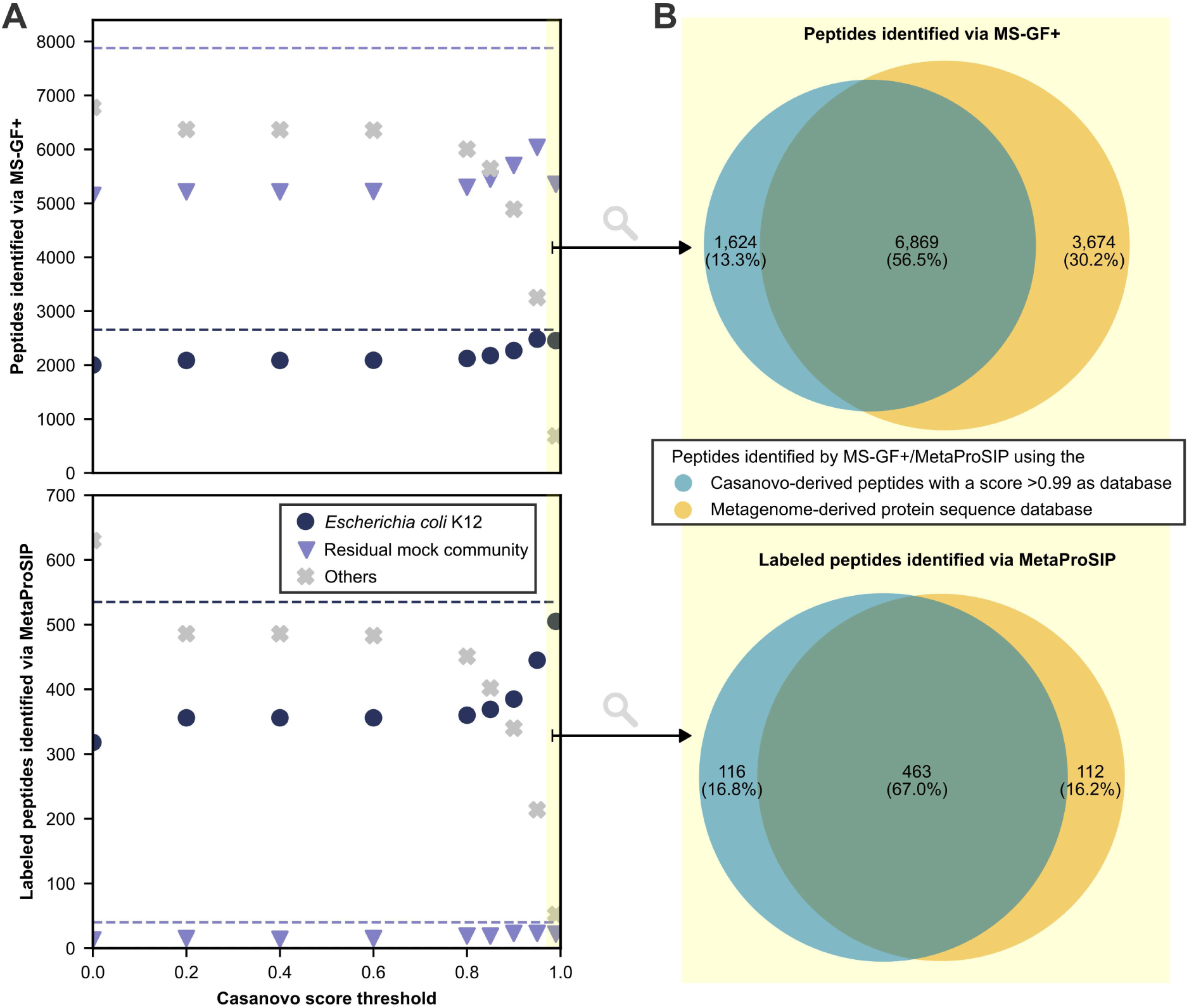
Comparison of unlabeled and ¹³C-labeled peptide identifications in a mock microbial community spiked with ^13^C-labeled *Escherichia coli* K12 cells using MS-GF+ and MetaProSIP with *de novo* peptide databases (1% PSM-level FDR) and a metagenome-derived protein database (2% PSM-level FDR). Raw data was obtained from [27]. The score threshold of >0.99 is highlighted in yellow. Only peptides detected in at least two samples are plotted. **A**: *De novo* peptide databases were generated using Casanovo and filtered at varying quality score thresholds. Identified peptides are color-coded based on their presence in the *E. coli* K12 reference protein database (PRIDE accession number: PXD024285) or common contaminants (navy circles), the metagenome-derived database of the residual mock community (PRIDE accession number: PXD006118, purple triangles), or neither (grey crosses). Dashed lines represent identifications from the metagenome-derived database. **B**: Area-proportional Venn diagram of peptides identified via MS-GF+ and MetaProSIP using either the metagenome-derived database or a *de novo* database filtered at a Casanovo quality score >0.99.

Casanovo-derived databases outperformed PepNet-derived ones by identifying more peptides from expected community members and fewer unrelated sequences (Figure 3A and Supplementary Figure S1). The Casanovo-derived database, filtered by a quality score >0.99 (CS>0.99), contained 14,700 peptides and led to the highest number of labeled *E. coli* K12 peptide identifications among the *de novo* databases. To assess the proportion of false discoveries within the set of identified peptides, we applied an entrapment approach. With 1% PSM-level FDR, the estimated false discovery proportion was 1.15% for the CS>0.99 database and 0.71% for the metagenome-derived database. Increasing the PSM-level FDR threshold for the metagenome-derived database to 2% yielded a comparable false discovery proportion of 1.19%. We then compared peptide identifications across the CS>0.99 database (with 1% PSM-level FDR) and the metagenome-derived databases (with 2% PSM-level FDR) (Figure 3B). Most unlabeled and labeled peptides were identified by both databases. Each database also contributed unique identifications, with the CS>0.99 database identifying slightly more unique labeled peptides, and the metagenome-derived database yielding more unique unlabeled peptides.

To test why the number of identified labeled *E. coli* K12 peptides increases with more stringent database filtering, we manually investigated the gain and loss of peptide spectrum matches between databases filtered with different quality scores. We observed two opposing effects driven by filtering based on Casanovo quality scores: First, *E. coli* K12 peptides with Casanovo scores below the threshold are excluded and thus cannot be identified, leading to a loss in *E. coli* K12 identifications. Second, isobaric non-*E. coli* K12 peptides with Casanovo scores below the threshold are excluded and thus cannot compete with *E. coli* K12 peptides for the same spectrum during the database search, increasing the number of identified *E. coli* K12 peptides. For example, comparing databases filtered at different Casanovo score thresholds (CS>0.95 and CS>0.99), the peptide “AAAFEGELLPA<u>SQL</u>DR” (*E. coli* K12) is present in both databases, while the isobaric variant “AAAFEGELLPA<u>KAE</u>DR” (non-*E. coli* K12) appears only in the CS>0.95 database. When both peptides are present in the database (CS>0.95), MS-GF+ assigns the spectrum to the non-*E. coli* K12 peptide based on a better SpecEvalue. When the isobaric non-*E. coli* K12 variant is absent from the database (CS>0.99), the same spectrum is matched to the *E. coli* K12 sequence, which is classified as labeled in both cases. However, only the match to the *E. coli* K12 sequence contributes to labeled *E. coli* K12 peptide identifications. Importantly, when the number of labeled *E. coli* K12 peptide identifications increases under more stringent filtering, this indicates that the second effect—removal of competing isobaric non-*E. coli* K12 variants—outweighs the first effect—loss of *E. coli* K12 peptides due to filtering. Indeed, when comparing detected labeled peptides between using the CS>0.95 database and the CS>0.99 database, 35 labeled *E. coli* K12 peptides were lost with more stringent filtering (effect 1), while 95 labeled peptides were gained due to the removal of competing isobaric non-*E. coli* K12 variants (effect 2). Since *E. coli* K12 was isotopically labeled in the experiment, we expect true *E. coli* K12 peptides to be labeled. Therefore, labeled *E. coli* K12 identifications carry higher confidence than unlabeled ones, and an increase in labeled *E. coli* K12 peptides suggests improved identification specificity. For this reason, we selected CS>0.99 as the default database for downstream analyses (Figure 4).

**Figure 4:**
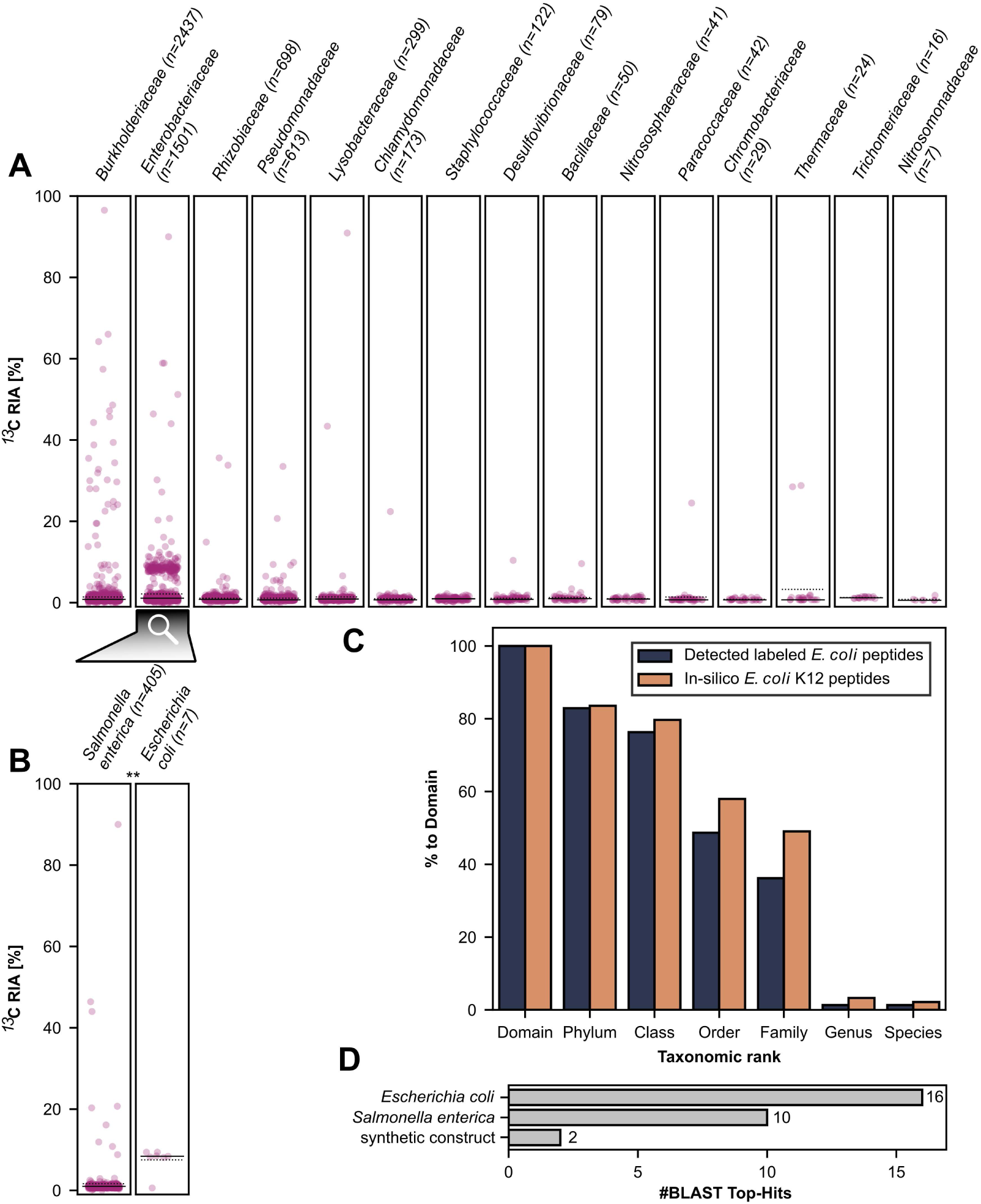
Analysis of ^13^C relative isotope incorporation (RIA) into peptides of different taxa identified in a mock microbial community with spike-in of ^13^C-labeled *E. coli* cells. RIA values are shown for peptides detected in at least two replicates and for taxa for which at least three peptides were detected. Dots indicate peptides assigned to (**A**) the detected families or (**B**) the detected *Enterobacteriaceae* spp.. Not all *Enterobacteriaceae* peptides were species-specific. ‘n’ denotes the number of peptides identified for a taxon across all replicates. Solid lines indicate medians. Dotted lines indicate means. Statistically significant differences in RIA between species are marked with ‘**’ based on Student’s t-test for the means of two independent samples with p < 0.01. **C**: Comparison of drop-off rates from *in silico* digestion of the *E. coli* K12 proteome (1,344 domain-specific peptides) and labeled peptides matching the lineage of *E. coli* detected in at least two replicates of the mock microbial community with spike-in of ^13^C-labeled *E. coli* cells (190 domain-specific peptides). **D**: Number of best hits when running BLASTP on the 47 labeled peptides not assigned to any lowest common ancestor by Unipept and detected in at least two replicates of the mock microbial community with spike-in of ^13^C-labeled *E. coli* cells. Sequence similarity search was performed using BLASTP against the nr database. Organisms with only one hit and hits with e-value ≥ 0.001 were removed before plotting.

Using the CS>0.99 database, we investigated the level of taxonomic resolution that could be identified for the labeled organisms within the unlabeled residual community. For this purpose, we inferred the taxonomy of all identified peptides with a lowest-common-ancestor approach using Unipept and plotted the peptides with their assigned family against the calculated ^13^C RIA (Figure 4A). From the 8,493 peptides consistently detected with the MetaProSIP workflow in at least two replicates, 7,934 were assigned to a lowest common ancestor using Unipept. Additionally, 579 of the 8,493 peptides were labeled in at least two sample out of which 532 were assigned to the lowest common ancestor via Unipept.

After filtering for peptides detected in at least two replicates and for taxa for which at least three peptides were detected, we identified all non-viral orders in the mock community, 14 out of the 16 families, 15 out of the 20 genera, 15 out of the 23 species expected in the mock community (Supplementary Table S2). We did not detect any viruses present in the mock community. However, we detected two unexpected taxa: 6 fungal peptides assigned to *Knufia peltigerae* (family *Trichomeriaceae*) and 4 peptides assigned to *Chromobacterium sphagni*. Further analysis revealed that these peptides also match *Stenotrophomonas* spp. (NCBI: WP_025877809.1) and other *Chromobacterium* spp., including *Chromobacterium violaceum* (NCBI: WP_076225604.1, WP_101707107.1, and GCF_019460125.1), which are known members of the mock community, suggesting misassignment by Unipept’s lowest common ancestor algorithm.

Most of the identified peptides cluster around an RIA of 1.1%, corresponding to the natural ^13^C-abundance indicative of unlabeled peptides. Only for *Enterobacteriaceae* we observed a second cluster of peptides centered between an RIA of approximately 7% and 11%. *Salmonella enterica* and *E. coli* were the only members of *Enterobacteriaceae* identified at higher taxonomic resolution, whereby peptides assigned to *E. coli* exhibited a median RIA of 8.4%, which is significantly higher (p < 0.01) than that found for peptides from *S. enterica,* indicating that only *E. coli* peptides were labeled (Figure 4B).

To test for hidden labeled populations, we applied three approaches. First, we compared the number of labeled peptides assigned to *E coli* or superordinate taxa of its lineage with the number of labeled peptides assigned to other taxa. Our results showed that most labeled peptides were assigned to *E. coli* or superordinate taxa of its lineage, indicating a high population share of *E. coli* in the labeled subpopulation of the sample (Supplementary Table S3). Secondly, we compared the theoretical drop-off rate of *E. coli* K12 peptides with the observed drop-off rate of detected labeled peptides. Drop-off rates describe the decline in taxon-specific peptide counts from more general to more specific taxonomic levels [51]. By comparing observed and theoretical drop-off rates, it is possible to assess whether peptide counts at a given taxonomic level can account for counts at more general levels [51]. Consequently, drop-off rates can reveal hidden subpopulations not captured by the taxonomic database. Here, the observed drop-off rate deviated only slightly from the theoretical one, indicating the absence of hidden side populations (Figure 4C). Thirdly, we performed a sequence similarity search for the 47 labeled peptides that Unipept could not assign to any lowest common ancestor (Figure 4D). For 22 of those peptides, BLASTP found highly similar sequences with an e-value < 0.001. After removing organisms with only one hit, all highly similar sequences were related to *E. coli*, its close relative *S. enterica* [58], or synthetic constructs. Consequently, all performed tests indicate the absence of hidden labeled subpopulations in accordance with the setup of the defined microbial community.

### Case study 1: Investigating trophic relations in a continuous flow reactor for the cultivation of anammox bacteria by tracing ^13^C incorporation into peptides

To test the power and applicability of our approach on real samples, we analyzed proteomics data from a continuous flow anammox reactor enriched with ‘*Candidatus* Kuenenia stuttgartiensis’ strain CSTR1 at three different time points after changing the carbon source from non-labeled to ^13^C-labeled bicarbonate while leaving nitrogen sources unchanged.

Again, it was possible to detect and subsequently quantify the RIA of peptides using *de novo* peptide databases. The *de novo* peptide databases had a size of 5,313 to 303,595 non-redundant peptides (Supplementary Table S1). For comparison, the genome-derived protein sequence database was constructed from 4,963 protein-coding genes translated from the complete genome of ‘*Ca.* Kuenenia stuttgartiensis’ strain CSTR1 (NCBI accession: CP049055.1) and appended with universal contaminant sequences [59, 60]. For samples acquired on day 21 (5.25 × hydraulic retention time), it was necessary to mix labeled peptides with unlabeled peptides from day 0 to enable *de novo* peptide sequencing and subsequent RIA calculation by MetaProSIP (Supplementary Table S4). Across all samples, we detected more labeled ‘*Ca.* Kuenenia stuttgartiensis’ strain CSTR1 peptides and less non-CSTR1 labeled peptides with the Casanovo-derived databases compared to the PepNet-derived ones (Supplementary Figure S2). Similarly, we compared different quality score thresholds for filtering *de novo* identified peptides before creating the database for SIP analysis (Figure 5A).

**Figure 5:**
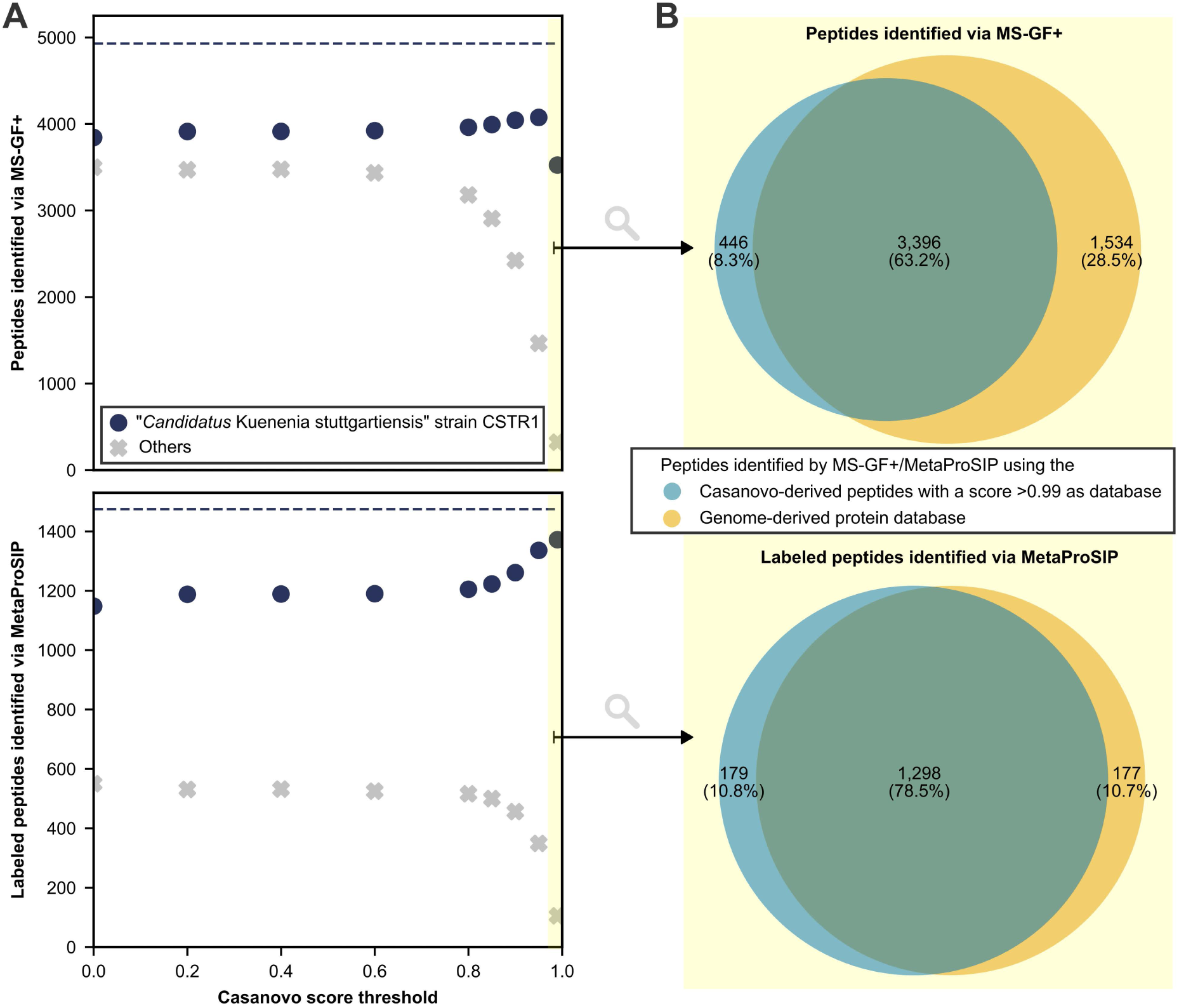
Comparison of unlabeled and ¹³C-labeled peptide identifications using MS-GF+ and MetaProSIP in an enrichment of ‘*Candidatus* Kuenenia stuttgartiensis’ strain CSTR1 from a continuous flow reactor fed with ^13^C-bicarbonate with *de novo* peptide databases (1% PSM-level FDR) and a genome-derived protein database (2% PSM-level FDR). The score threshold of >0.99 is highlighted in yellow. Only peptides detected in at least two samples are plotted. **A**: *De novo* peptide databases were generated using Casanovo and filtered at varying quality score thresholds. Navy circles represent peptide identifications matching the genome-derived database (NCBI: CP049055.1) or universal contaminants; grey crosses represent unmatched peptides. Dashed lines show identification counts using the genome-derived database. **B**: Area-proportional Venn diagram of peptides identified via MS-GF+ and MetaProSIP using either the genome-derived database or a *de novo* database filtered at a Casanovo quality score >0.99 [61].

Using the *de novo* peptide database filtered for a Casanovo quality score greater than 0.99 (CS>0.99), MS-GF+ identified 3,842 peptides present in at least two samples. Of these, 3,525 matched known proteins from ‘*Ca.* Kuenenia stuttgartiensis’ strain CSTR1 or universal contaminants. Subsequently, MetaProSIP identified 1,477 labeled peptides present in at least two samples, with 1,372 matching proteins from ‘*Ca.* Kuenenia stuttgartiensis’ strain CSTR1 or universal contaminants.

To estimate false discoveries, we applied an entrapment approach using both the CS>0.99 peptide database and the genome-derived database. At 1% PSM-level FDR, the estimated false discovery proportion for peptides detected in at least two samples was 1.09% using the CS>0.99 database and 0.64% using the genome-derived database. Increasing the FDR threshold for the genome-derived database to 2% yielded a more comparable false discovery estimate of 1.20%. With a 2% PSM-level FDR, the genome-derived database identified 4,930 peptides in at least two samples, including 1,475 labeled peptides. Comparison of peptide identifications (Figure 5B) showed substantial overlap between peptides identified with the CS>0.99 database and peptides identified with the genome-derived databases, with each approach also contributing unique peptides.

To investigate the ^13^C incorporation into taxa over time, we used the CS>0.99 database in MS-GF+ and MetaProSIP and analyzed the RIA and taxonomy of the 1,947 peptides consistently detected at each time point in at least two replicates (Figure 6). 1,846 of the 1,947 peptides, were successfully assigned to a lowest common ancestor using Unipept. Additionally, 1,290 of the 1,947 peptides were labeled in at least two samples out of which 1,225 were assigned to lowest common ancestor via Unipept. At the phylum level, most detected peptides were assigned to *Plancomycetota*, consistent with their high abundance in the reactor (Figure 6A). Some peptides were assigned to the phylum *Chordata*, indicating human contamination, and a few were linked to *Pseudomonadota*, previously described as a subpopulation of the reactor [62]. At the species level, we only detected ‘*Ca.* Kuenenia stuttgartiensis’ belonging to the phylum *Planctomycetota,* suggesting high enrichment of the culture (Figure 6B). Substantial incorporation of ^13^C has been detected only for peptides assigned to ‘*Ca.* Kuenenia stuttgartiensis’ or superordinate taxa of its lineage, indicating its activity and assimilation of ^13^C-bicarbonate (Supplementary Table S5). Specifically, we observed a significant (p < 0.001) increase of ^13^C RIA in ‘*Ca.* Kuenenia stuttgartiensis’ peptides at each investigated time point, resulting in a median RIA of 95.7% after 5.25 × hydraulic retention time demonstrating almost complete substitution of ^12^C with ^13^C.

**Figure 6:**
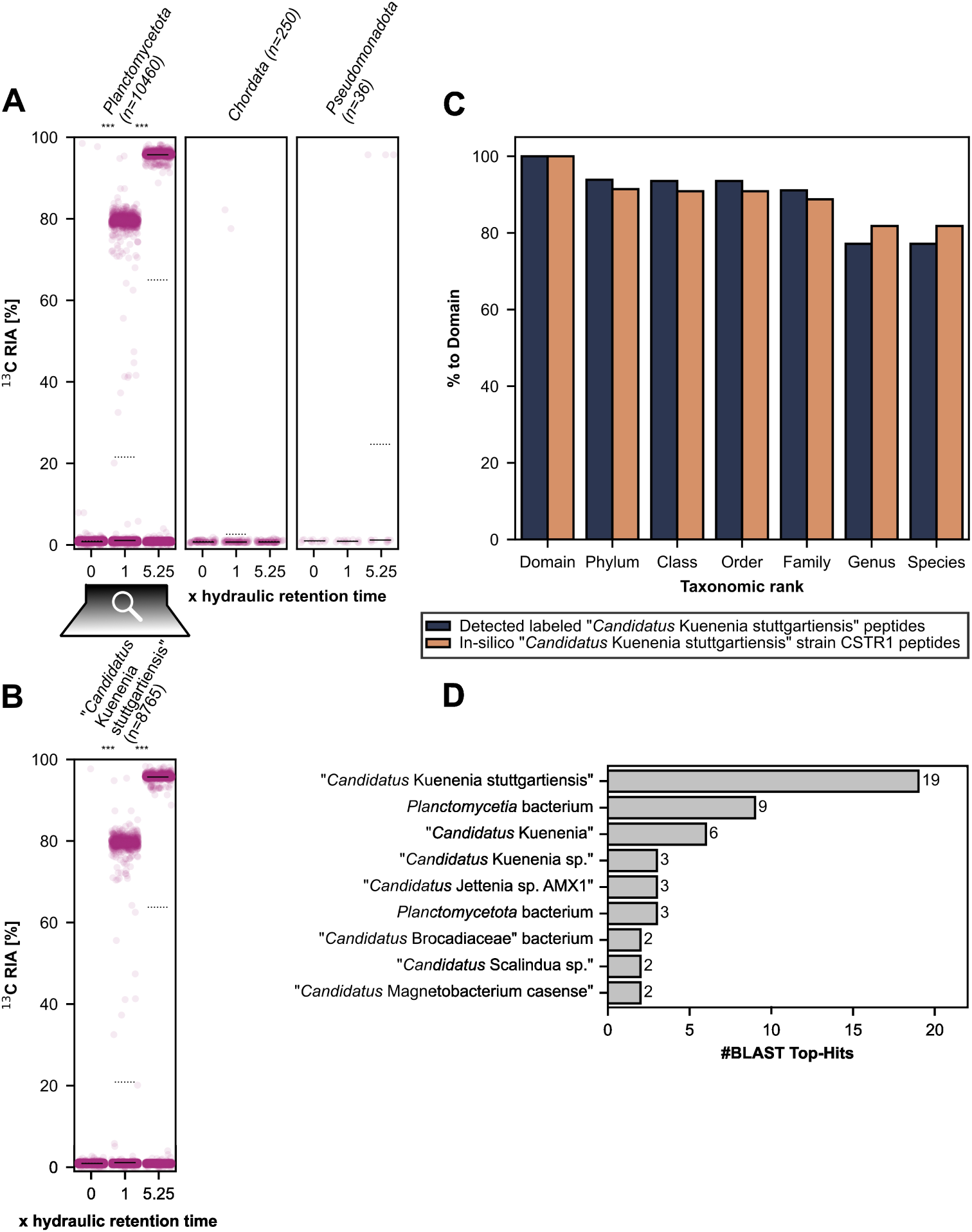
Distribution of the ^13^C relative isotope abundance (RIA) in detected peptides across taxa identified in samples from a continuous flow reactor enriching ‘*Candidatus* Kuenenia stuttgartiensis’ strain CSTR1 at different time points after switching to medium containing ^13^C-bicarbonate. Dots indicate peptides assigned to detected phyla (**A**) or species (**B**). RIA values are shown for peptides detected in at least two replicates at each time point. Taxa are shown for which at least three peptides were detected. ‘n’ denotes the number of peptides assigned to a taxon across all replicates and time points. Solid lines indicate medians. Dotted lines indicate means. Statistically significant differences in RIA between time points are indicated with ‘***’ based on Student’s t-test for the means of two independent samples with p < 0.001. **C**: Comparison of drop-off rates from *in silico* digestion of the ‘*Candidatus* Kuenenia stuttgartiensis’ strain CSTR1 proteome (2,544 domain-specific peptides) and labeled peptides matching the lineage of ‘*Candidatus* Kuenenia stuttgartiensis’ detected in at least two samples of the continuous flow reactor enriching ‘*Ca*. Kuenenia stuttgartiensis’ strain CSTR1 (894 domain-specific peptides). **D**: Top-hit species distribution after sequence similarity search of the 65 labeled peptides not assigned to any lowest common ancestor by Unipept but detected in at least two samples of the continuous flow reactor enriching ‘*Candidatus* Kuenenia stuttgartiensis’ strain CSTR1. The sequence similarity search was performed with BLASTP against the nr database. Organisms with only one hit and hits with e-value ≥ 0.001 were removed before plotting.

To investigate for labeled populations not covered by Unipept, we again compared observed and theoretical drop-off rates of peptides (Figure 6C). The drop-off rate of detected labeled peptides closely resembled the theoretical one, with a maximum deviation of only 6.0% (Supplementary Table S5). This suggests that all detected labeled peptides were derived from ‘*Ca.* Kuenenia stuttgartiensis’.

Furthermore, we searched for sequences similar to the 65 labeled peptides that Unipept could not assign to any taxon (Figure 6D). For 18 of those peptides, BLASTP identified highly similar sequences with e-value < 0.001. After removing organisms with only one hit, most of the highly similar sequences belonged to ‘*Ca.* Kuenenia stuttgartiensis’ or superordinate taxa of its lineage. In total five hits with high sequence similarity were affiliated with ‘*Candidatus* Jettenia sp. AMX1’ and ‘*Candidatus* Scalindua sp.’ which are anammox bacteria. In brief, our results indicate that ‘*Ca.* Kuenenia stuttgartiensis’ also dominates the labeled population in the reactor.

### Case study 2: Investigating species-specific and function-specific activity between two different simulated diets in a human distal gut microbiota model by tracing ^2^H and ^18^O incorporation into peptides

To test our strategy for comparative purposes and also for stable isotopes other than ^13^C, we analyzed previously published raw data (PRIDE accession number: PXD024291) from a model community of the human gut microbiota (‘Robogut community’), inoculated with 63 isolates, and cultivated for 12 h in a protein-rich or fiber-rich medium simulating different diets [17, 27]. The medium contained unlabeled water (for controls) or water in which 25% was substituted with either ^2^H2O or H2^18^O. First, we compared different *de novo* peptide databases for identifying unlabeled and labeled peptides expected to be present in the sample based on the published reference database. The *de novo* peptide databases contained 48,715 to 715,031 non-redundant peptide sequences (Supplementary Table S1). We identified more Robogut peptides with databases created using Casanovo compared to those created using PepNet, for both investigated stable isotopes ^18^O and ^2^H (Figure 7 and Supplementary Figure S3). The database for which peptides were *de novo* sequenced by Casanovo and filtered with a score threshold of 0.99 (CS>0.99) led to the most identifications of labeled Robogut peptides among all investigated databases. Furthermore, we identified more labeled peptides in samples cultivated with ^18^O than with ^2^H and more unlabeled peptides in the ^2^H samples. We identified more Robogut peptides with the CS>0.99 peptide database than with the genome-derived restricted reference protein database used in previous analyses of this dataset [17, 27]. The genome-derived restricted reference protein database was generated by Kleiner et al. [27] (PRIDE accession number: PXD024291) from UniProt reference proteomes of the species most closely related to the 63 isolates identified by 16*S* rRNA gene amplicon sequencing and used as inoculum by Starke et al. [17]. This database was appended with universal contaminant sequences [59], resulting in a total database size of 213,931 protein sequences.

**Figure 7:**
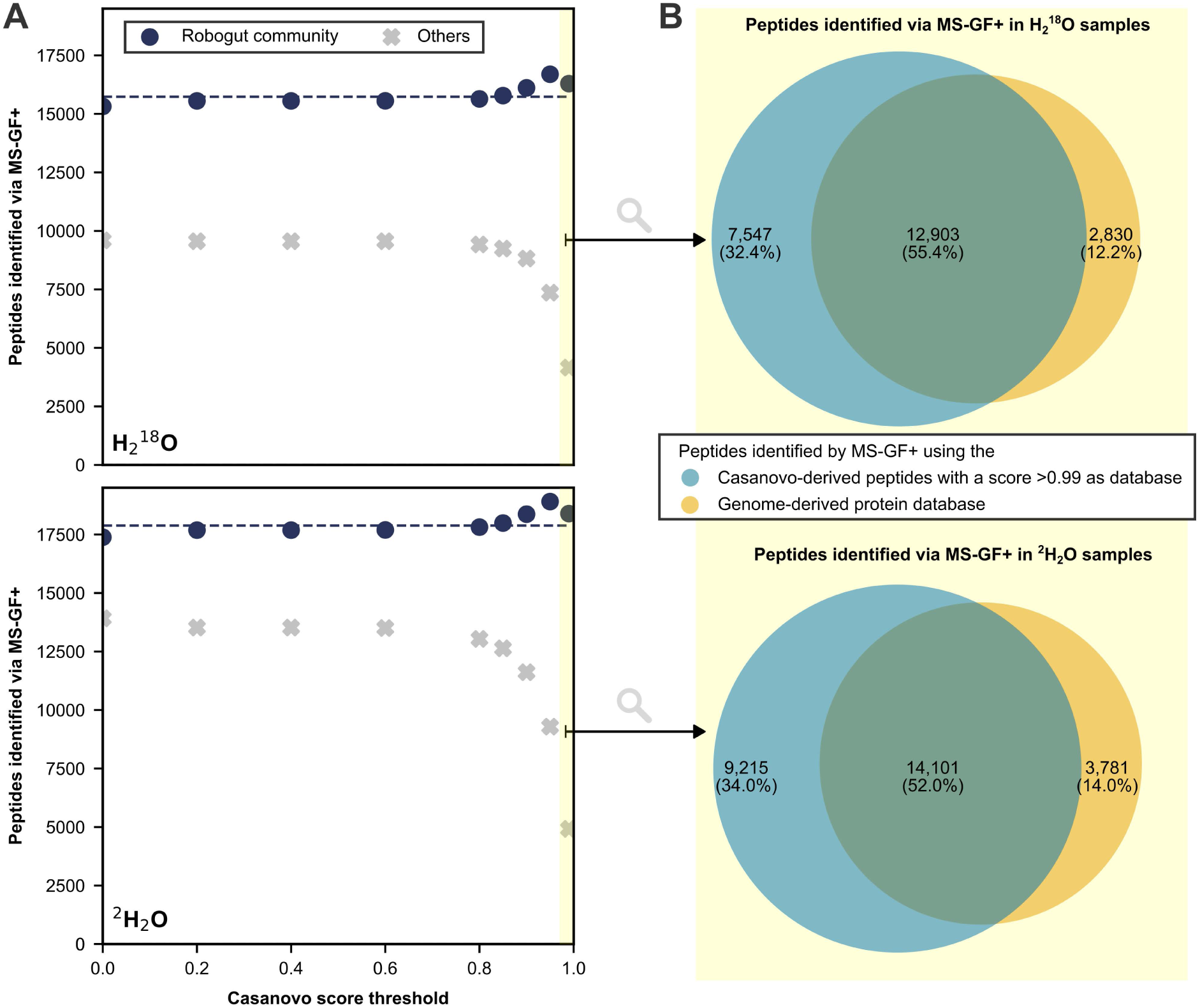
Comparison of *de novo* peptide databases for identifying peptides in a model of the human gut microbiota (Robogut community) cultivated with H_2_ ^18^O or ^2^H2O, respectively, and obtained from [17] via MS-GF+ (1% PSM-level FDR). The score threshold of >0.99 is highlighted in yellow. Only peptides detected in at least two samples are plotted. Only peptides detected in at least two samples are plotted. **A**: *De novo* peptide databases were assembled using Casanovo after filtering all identifications with defined thresholds for the quality score. Navy circles denote the number of identified peptides present in the genome-derived reference database or universal contaminants. Grey crosses denote the number of identified peptides absent in the genome-derived reference database and universal contaminants. Dashed lines represent the number of peptides identified using the genome-derived reference database appended with universal contaminants. **B**: Area-proportional Venn diagram of peptides identified via MS-GF+ using either the Casanovo-derived peptides with a score >0.99 for database search or the genome-derived protein sequence database for database search [61].

To assess the reliability of peptide identification, we estimated false discovery proportions using an entrapment approach across two databases: the CS>0.99 *de novo* peptide database and the genome-derived protein sequence database. The entrapment analysis estimated a false discovery proportion of 0.04% (for ^18^O samples) and 0.23% (for ^2^H samples) for peptides identified from the CS>0.99 database in at least two samples. The genome-derived protein sequence database showed a slightly higher, yet comparably low, false discovery proportion of 0.28% (for ^18^O samples) and 0.64% (for ^2^H samples) for peptides meeting the same criterion. We further evaluated the overlap between peptides identified by the two databases (Figure 7B). Again, most peptides were identified by both the CS>0.99 and genome-derived databases. Notably 32.4% to 34.0% of peptides were only identified when using the CS>0.99 database and 12.2% to 14.0% were exclusively identified with the genome-derived database.

Next, we analyzed the RIA and taxonomy of peptides identified under different dietary conditions using the CS>0.99 database. Using the CS>0.99 database, MS-GF+ identified 35,115 non-redundant peptides, out of which 8,099 and 8,999 peptides were detected in at least two replicates of both media compositions in presence of ^18^O and ^2^H, respectively. From these 8,099 and 8,999 peptides, 7,669 and 8,430 were assigned to a lowest common ancestor using Unipept, respectively.

Overall, the incorporation of ^18^O was substantially higher than the incorporation of ^2^H. Moreover, the identified peptides generally had significantly higher RIA (p < 0.01) in protein-rich compared to fiber-rich conditions (Supplementary Figure S4).

After inferring the taxonomy of peptides via Unipept, we identified 22 species present in all tested setups (Figure 8). Notably, 41 and 16 unique peptides assigned to *Copranaerobaculum intestinale* and *Hominenteromicrobium mulieris*, respectively, were reliably detected in at least 2 biological replicates of ^18^O- and ^2^H-labeled samples — both the peptides and the species were absent from the reference database used in previous analyses [17, 27]. The presence of *C. intestinale* and *H. mulieris* was validated through a separate database search against their reference proteomes (Uniprot: UP000434036 and UP001199424), identifying 110 and 133 *C. intestinale* peptides, as well as 285 and 329 *H. mulieris* peptides, absent from the reference database in at least two biological replicates of ^18^O- and ^2^H-labeled samples. In ^18^O incubated samples, more taxa were found to differ significantly in their RIA between the two simulated diets than ^2^H incubated samples. Specifically, we detected significantly higher RIA under high protein conditions compared to high fiber conditions for peptides assigned to *Akkermansia muciniphilia*, *Bacteroides ovatus*, *Bacteroides uniformis*, *Clostridium hylemonae*, *Clostridium symbiosum*, and *Copraanaerobaculum intestinale*, as well as significantly lower RIA for *Hominenteromicrobium mulieris* and *Hungatella hathewayi*.

**Figure 8:**
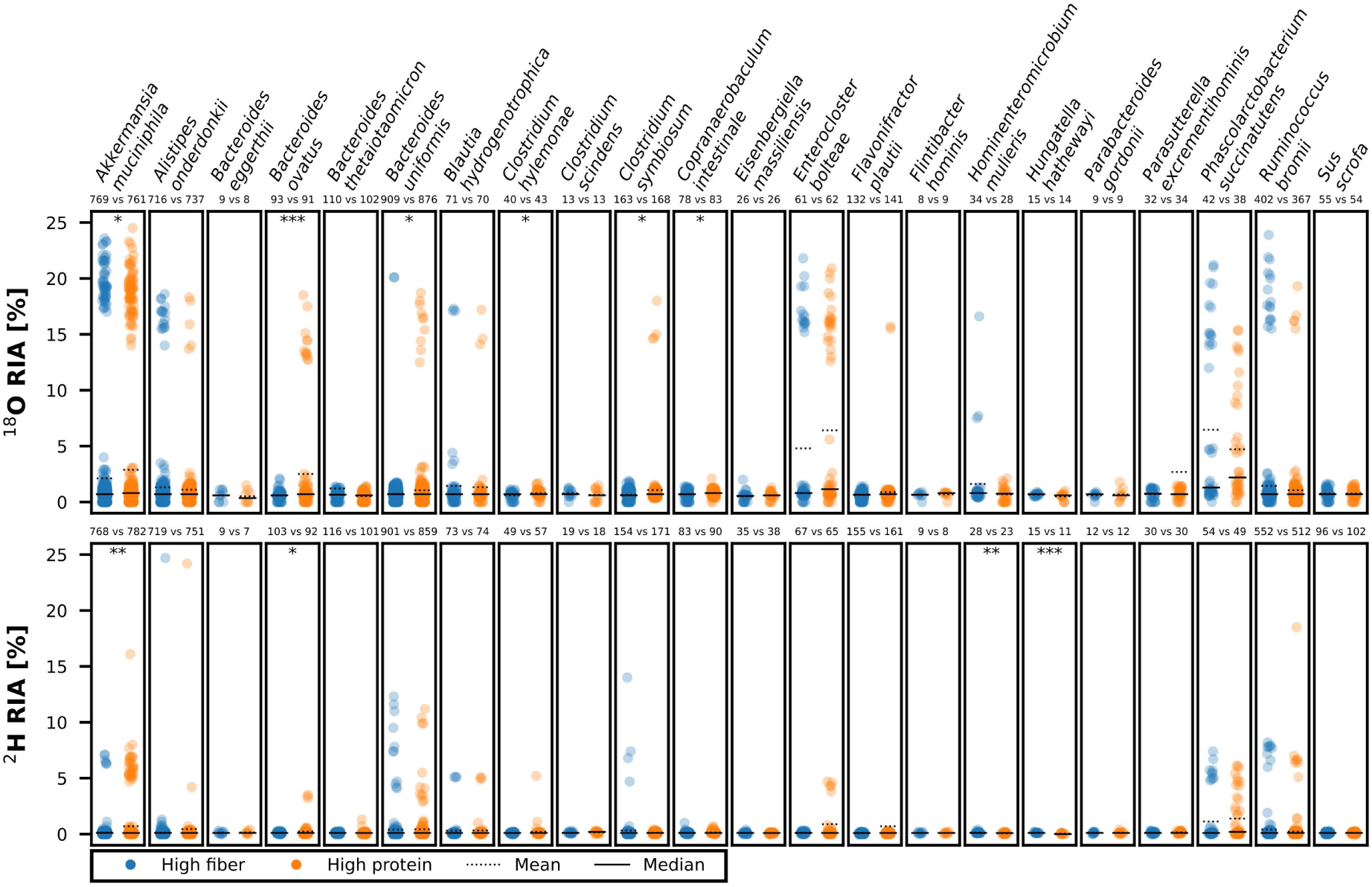
Distribution of the ^18^O and ^2^H relative isotope abundance (RIA) in detected peptides across species identified in a model of the human gut microbiota cultivated in either a high fiber or high protein media in the presence of either H_2_ ^18^O or ^2^H2O. RIA values are shown for peptides detected in at least two replicates of both media compositions. Species detected in the presence of H_2_ ^18^O and ^2^H2O are plotted for which at least three peptides were detected. ‘n’ denotes the number of peptides assigned to a specific taxon across triplicates. Statistically significant differences in RIA between media are indicated with ‘*’, ‘**’, and ‘***’ based on Student’s t-test for the means of two independent samples with p < 0.05, p < 0.01, and p < 0.001, respectively. To improve readability, outliers were removed from the raw graph (Supplementary Figure S5).

To assess changes in activity related to individual biological processes, we assigned identified peptides and their RIA to biological processes via gene ontology (GO) numbers using Unipept. For an overview, GO numbers were first merged into GO slims of the prokaryote subset (Figure 9).

**Figure 9:**
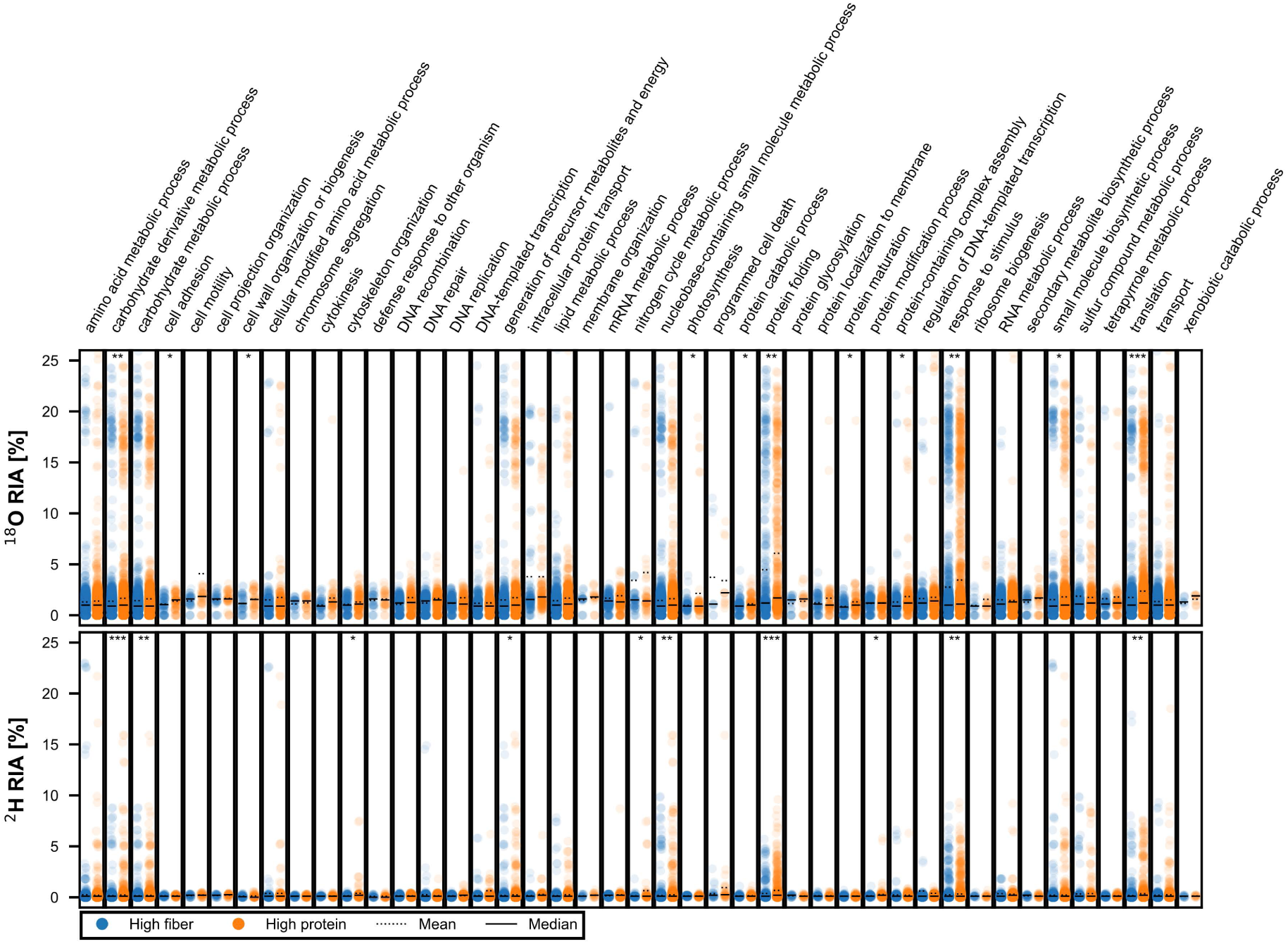
Distribution of the ^18^O and ^2^H relative isotope abundance (RIA) in detected peptides across general biological processes of prokaryotes (goslim_prokaryote) identified in a model of the human distal gut microbiota cultivated in either a high fiber or high protein medium in the presence of either H2^18^O or ^2^H2O. RIA values are shown for peptides detected in at least two replicates of both media and for biological processes detected in presence of H2^18^O and ^2^H2O for which at least three peptides were detected (10 to 3457 peptides per strip). Statistically significant differences in RIA between media are indicated with ‘*’, ‘**’, and ‘***’ based on Student’s t-test for the means of two independent samples with p < 0.05, p < 0.01, and p < 0.001, respectively. To facilitate readability, outliers were removed from the raw graph (Supplementary Figure S6).

For most biological processes, peptides exhibited similar RIA values between the diets, while they exhibited significantly different RIAs for other processes. Specifically, peptides assigned to carbohydrate derivative metabolic processes, protein folding, response to stimulus, and translation exhibited significantly higher ^2^H and ^18^O RIAs in the high protein medium. Peptides assigned to cell adhesion, cell wall organization or biogenesis, photosynthesis, protein catabolic processes, protein maturation, protein-containing complex assembly, and small molecule biosynthetic processes only exhibited significantly higher ^18^O RIA in the protein-rich medium. Peptides assigned to carbohydrate metabolic processes, cytoskeleton organization, generation of precursor metabolites and energy, nitrogen cycle metabolic processes, nucleobase-containing small molecule metabolic processes, and protein modification processes exhibited significantly higher ^2^H RIA in the protein-rich medium.

To further investigate biological processes related to dietary differences, we specifically analyzed all detected GO child terms of the parental GO slims of amino acid metabolic processes (GO:0006520), carbohydrate derivative metabolic processes (GO:1901135), and carbohydrate metabolic processes (GO:0005975). The identified peptides were assigned to a total of 160 biological processes related to the above-mentioned GO slims, out of which 28 biological processes exhibited significant differences (p < 0.05) in ^2^H or ^18^O RIA between both media compositions (for specific processes see Supplementary Table S6).

Differences in RIA related to amino acid or carbohydrate metabolism were isotope-specific. In the protein-rich medium, significantly higher ^18^O and ^2^H RIAs were observed in peptides associated with L-histidine catabolism to glutamate and formate, as well as carbohydrate metabolism. The protein-rich medium also showed significantly elevated ^18^O RIA in peptides linked to acetylglucosamine, arabinose, and fucose catabolisms, as well as fructose-6-phosphate metabolism, while ^2^H RIA was significantly higher in peptides involved in aminoacylation, biosynthesis of arginine, UTP, and GMP, as well as aspartate, glucose, and fructose-1,6-bisphosphate metabolism. Conversely, the fiber-rich medium exhibited significantly higher ^18^O RIA in peptides related to amino acid biosynthesis (particularly valine and isoleucine), and higher ^2^H RIA in peptides associated with cellulose catabolism, as well as arabinose and nucleoside metabolisms.

## Discussion

Our results show that *de novo* peptide sequencing can generate sample-matching databases for ^2^H, ^13^C, and ^18^O Protein-SIP analyses without prior knowledge of community compositions. In our work, databases derived from Casanovo allowed for the identification of consistently more peptides expected in the sample than PepNet-derived databases. Filtering *de novo* identified peptides to those with a quality score >0.99 before assembling the database (CS>0.99) enabled the identification of more labeled peptides expected in the sample than using a non-filtered database, supporting the conclusion that inflated database size can challenge database search algorithms, as reported previously [44, 48]. Previous studies have implemented hybrid approaches where *de novo* identified peptides are subsequently used for database searching, but did not validate the FDR [56, 57]. Our study further advanced these approaches by validating the FDR via an entrapment strategy, optimizing quality score thresholds, and integrating it with stable isotope probing.

At a comparable false discovery proportion of around 1% at the peptide level, the CS>0.99 databases enabled the identification of 77.9–130.4% of the number of unlabeled peptides and 100.1–112.0% of the number of the labeled peptides relative to the (meta)genome-derived databases. A substantial proportion of peptides, 52.0–63.2% of unlabeled and 67.0–78.5% of labeled peptides, were consistently identified by both database types, indicating high reliability. Notably, 8.3–34.0% of unlabeled and 10.8–22.0% of labeled peptides were uniquely identified using the CS>0.99 databases, highlighting their ability to detect peptides not captured by (meta)genome-derived databases. Such peptides could originate (1) from sequences that are not included in the metagenome bins, (2) from species not covered by the sequenced metagenomes (if only a portion of samples were sequenced for metagenome, which is usually the case), or (3) from contaminating organisms/proteins introduced during sampling and sample treatment. These results indicate that while CS>0.99 databases perform comparably to (meta)genome-derived databases in terms of peptide identifications, they can also provide complementary peptide coverage, particularly for labeled peptides. Thus, using metagenome-derived protein sequence databases in combination with *de novo* peptide databases may enhance sensitivity and improve the overall depth of protein-SIP analyses.

Based on the analyzed data and the obtained results, we propose a modular Denovo-SIP pipeline (Figure 1). In the proposed Denovo-SIP pipeline, peptides are first *de novo* sequenced using Casanovo [54] to create a peptide sequence database for subsequent database-dependent search. The database-dependent search is carried out by MS-GF+ [63] and MetaProSIP [31] to validate peptides with a target-decoy approach and infer isotope incorporation. Unipept [49] and BLASTP [50] can then retrieve taxa and functions of peptides. Python scripts from the Denovo-SIP pipeline allow the integration of isotope incorporation data with functional and taxonomic annotation for subsequent analysis. The modular structure of the Denovo-SIP pipeline allows flexibility, enabling the integration of different algorithms and the replacement or supplementation of modules with alternative tools to meet specific needs. For example, Sage could increase the speed of database search [64]; Sipros 4 could improve the accuracy and sensitivity in detecting labeled peptides [32]; Calis-p could enhance the analysis of low amounts of isotope incorporation [27]; NovoLign could automatize sequence alignment of peptides [65]; and MetaX could further analyze peptide-associated taxa and functions [66]. Furthermore, *de novo* peptide sequencing algorithms are developing fast: algorithms such as π-HelixNovo [67], PowerNovo [68], and NovoB [69], recently reported to outperform Casanovo on some datasets, could be used in the Denovo-SIP pipeline. Moreover, rescoring algorithms could be employed to increase sensitivity and precision of *de novo* peptide sequencing [70] as well as database searches [71].

We first tested the Denovo-SIP pipeline by analyzing proteomic data from a defined microbial community containing ^13^C-labeled *E. coli* cells. Using the pipeline, we successfully identified the isotope labeling in *E. coli* and showed that the residual community was unlabeled. Notably, we were able to confidently identify *E. coli* K12 at the species level, which is particularly challenging because this strain shares a higher average nucleotide identity with the type strain of *Shigella flexneri* than with the type strain of *E. coli* [72]. We did not detect any viruses in the mock community and were unable to resolve the species of eight additional members of the mock community, which may have been caused by the raw data quality. Even in the original study using a genome-derived reference protein database, these species typically yielded only 0 to 5 peptide identifications per replicate (Supplementary Table S2) [27], supporting previous observations that data quality remains a major limitation in *de novo* peptide sequencing [53, 73]. Similarly, another bottom-up proteomics study reported difficulties in detecting viruses with low relative abundances in complex microbial communities [74]. Importantly, although some species could not be resolved, our pipeline successfully identified all non-viral members at the order level and 14 out of 16 families. This reinforces the importance of assessing microbial community compositions at multiple taxonomic levels, particularly when relying on lowest common ancestor assignments of peptides [51].

By using the Denovo-SIP pipeline, we could also confirm that the anammox bacterium ‘*Candidatus* Kuenenia stuttgartiensis’ strain CSTR1 used bicarbonate as its sole carbon source. Furthermore, we did not detect any other organisms in the reactor assimilating bicarbonate. This experiment demonstrated that Denovo-SIP can detect highly enriched peptides with RIA of more than 95% as well as differentially labeled peptides between time points. Therefore, Denovo-SIP can be applied to time-series studies.

We also employed the Denovo-SIP workflow to analyze previously published data from the human distal gut model microbiota cultivated in a fiber-rich or protein-rich medium containing either ^2^H- or ^18^O-labeled water. This approach is used to detect general activity as deuterium and oxygen from water is always incorporated when a microorganism is active. Through this analysis, we showed that changes in activity due to dietary compositions are species- and process-specific in agreement with previous analyses [17, 27]. Previous work reported that a fiber-rich medium increases protein abundance related to amino acid biosynthesis [17]. Our work specifically suggests that the biosynthesis of valine and isoleucine, as well as the catabolism of histidine and the metabolism of carbohydrates were affected by the medium composition. Lower ^2^H RIAs compared to ^18^O RIAs can be explained by abiotic hydrogen–deuterium exchange during sample preparation in unlabeled water, which causes partial loss of deuterium from exchangeable sites, whereas ^18^O is more stably retained [17, 75].

In conclusion, our results show that *de novo* peptide databases perform comparably to genome-derived databases in detecting labeled peptides. The main advantages of *de novo* peptides databases include saving time and costs associated with metagenome sequencing, reducing computational time during database search compared to genome-derived protein databases (Supplementary Table S7), and enabling the detection of foreign peptides, which are typically absent in reference databases. Such foreign peptides could originate from contamination [59], mutation [76], alternative splicing [77], or nonspecific cleavage [78]. Moreover, *de novo* peptide databases allow the detection of peptides from organisms not expected in the sample, as shown here for *C. intestinale* and *H. mulieris* in the human distal gut model. *C. intestinale* and *H. mulieris* were first described in 2022, after publication of the dataset in 2020. Thus, it is reasonable that *C. intestinale* and *H. mulieris* were absent in the published reference database [79, 80]. By this, we show that new insights can be gained by employing *de novo* sequencing for reanalyzing previously published datasets.

*De novo* peptide databases also have certain limitations for stable isotope probing. First, *de novo* sequencing was only successful on peptides with RIA of up to 5%. Usually such low or unlabeled peptides are present in the beginning or in controls of experiments allowing database creation for subsequent analyses of labeled samples. This limitation is not specific to *de novo* sequencing. To date, all major SIP tools (SIPPER [30], MetaProSIP [31], and Sipros [32]) require unlabeled reference samples to identify labeled peptides. In the rare case where no unlabeled peptides are available, biases due to undetected highly labeled peptides may occur and would require alternative analytical strategies. Secondly, *de novo* peptide sequencing is computationally intensive and often requires high-performance computing graphics processing units (GPUs) to achieve maximum performance. However, many algorithms, including Casanovo, remain usable via CPU on standard desktop systems, albeit with longer runtimes. Furthermore, the increasing availability of public cloud computing platforms (e.g., Galaxy, Google Colab) has made GPU-based *de novo* sequencing more accessible to the broader research community. Thirdly, *de novo* peptide databases contain amino acid sequences derived from mass spectra. Amino acid sequences are less variable than nucleic acid sequences. Hence, *de novo* peptide databases inherently result in lower taxonomic resolution than databases derived from nucleic acid sequences, such as metagenome sequencing. Additionally, we employed a peptide-centric approach, which avoids the protein inference problem but inherently results in lower taxonomic and functional resolution than protein-centric methods due to the shorter sequence length and limited specificity of peptides compared to proteins [81]. Furthermore, taxonomy inference via a lowest common ancestor approach relies on detecting taxon-specific peptides. The number of taxon-specific peptides decreases with increasing taxonomic resolution [82]. Moreover, the abundance of taxon-specific peptides varies between taxa [46]. Thus, the overall sensitivity and taxonomic resolution of lowest common ancestor approaches is dependent on the specific taxa present and their abundance. Still, we demonstrated that *de novo* peptide databases can achieve species-specific resolution when sufficient species-specific peptides are detected. Another limitation is that although identifying peptides by *de novo* sequencing does not require their presence in databases, interpreting the identified peptides by inferring proteins, taxa, or functions still requires a reference database. We propose using Unipept for taxa/function inference. Recently, Unipept update 6.0 has been announced to allow the analysis of non-tryptic peptides enhancing the synergy with *de novo* sequencing [83]. To get information about peptides that do not perfectly match, taxonomic drop-off rates and BLAST can be integrated into our automated workflow. In doing so, Denovo-SIP can assess the potential of unreferenced organisms. Estimating false-discovery rates of *de novo*-identified peptides is challenging [84], and currently no tools for *de novo* sequencing of labeled peptides exist. Therefore, our approach of assembling databases through *de novo* peptide sequencing – followed by database searches to verify these peptides and calculate isotope incorporation rates – shows promise. Denovo-SIP could advance the application of Protein-SIP across various microbiological research fields, serving as a complement to analyses based on (meta)genome-derived protein sequence databases or as a substitute in exploratory studies or contexts with limited resources.

## Methods

### Cultivation and protein extraction

Proteomics data of the anammox community were obtained with samples taken from a continuous stirred-tank reactor cultivating a mixed culture containing predominantly the anammox bacterium ‘*Candidatus* Kuenenia stuttgartiensis’ strain CSTR1 [62]. The reactor had an effective volume of 500 mL and was fed with two medium solutions containing 60 mM ammonium and 60 mM nitrite (concentration after mixing with ratio 1:1), respectively. Sodium bicarbonate (10 mM, concentration after mixing) was the only carbon source provided, provided with the nitrite medium. For a detailed medium recipe, please refer to [62]. The reactor was running stably with non-labeled bicarbonate at 30 °C, constant stirring, and 4-day hydraulic retention time. The cell density was around 2.5 × 10^8^ cells mL^-1^. At day 0, the nitrite medium containing non-labeled sodium bicarbonate was replaced with a medium containing ^13^C-labeled sodium bicarbonate, while all other medium components and reactor operational conditions remained the same. Three samples (as triplicates) were taken from the reactor on day 0 (non-labeled), day 4 (1 × hydraulic retention time), and day 21 (5.25 × hydraulic retention time). Samples of 1.5 mL were centrifuged at 10,000 × *g*, 10 °C for 10 min, and the pellets were frozen at -20 °C until proteomic analysis was done.

### Sample preparation and mass spectrometry

Saved pellets from reactor liquid were resuspended in 50 µL of 100 mM ammonium bicarbonate buffer, and went for a 3× freeze/thaw cycle to disrupt the cells (-80 °C followed by 1 min gentle shaking at 40 °C). Two microliter of 40 ng µL^-1^ bovine serum albumin was added to each sample as internal standard. Then, an equal volume of 10% (w/v) sodium deoxycholate (DOC) was added to each sample, before 12 mM dithiothreitol (DTT) and 40 mM 2-iodoacetamide (IAA) (both were final concentrations) were added sequentially to reduce and alkylate the disulfide bond (incubation conditions: 30 min, 37 °C on a rotary shaker at 400 rpm for DTT; and 45 min, room temperature, 400 rpm, in the dark for IAA). Afterwards, 100 mM ammonium bicarbonate was added to dilute DOC to 1% (w/v). Sequencing-grade trypsin (6.3 µL, 0.1 µg µL^-1^, Promega, Madison, USA) was then added to digest the sample at 37 °C, 400 rpm overnight. After trypsin digestion, neat formic acid was added to a final concentration of 2% (v/v), samples were vortexed, and two rounds of centrifugation at 16,000 × *g* for 10 min were performed to completely remove precipitated DOC. Peptides in the supernatant were desalted using 100 µL C18 tips (Pierce, Thermo Fisher Scientific, Massachusetts, USA) following the manufacturer’s instruction. Eluted peptides were vacuum dried and resuspended in 50 µL of 0.1% (v/v) formic acid. Peptide concentrations were determined based on the UV absorbance at 205 nm measured on a DS-11 micro-volume spectrophotometer (DeNovix, Delaware, USA). Peptides from day 4 (1 × hydraulic retention time) and day 21 (5.25 × hydraulic retention time) were mixed with peptides from day 0 in a 1:1 ratio before 3 µL were injected for measurement on nano-LC-MS/MS (Orbitrap) as described previously [60] except that the MS2 resolution was lowered from 60,000 to 30,000. The proteomics data were deposited at ProteomeXchange via the PRIDE partner repository under the accession number PXD057215 (reviewer token: ejwbd9cUNtVw).

### Generation of *de novo* peptide databases

Mass spectrometric raw files (Supplementary Table S8) were converted into mgf format after peak-picking, removing zero values in spectra, encoding in 64-bit precision, and zlib compression using msConvert (ProteoWizard v3.0.23304) [85]. Mass spectra were translated to peptide sequence lists by *de novo* sequencing using PepNet (v0.3.2) [55] and Casanovo (v3.5.0) [54] with the pre-trained models. Peptide sequence lists were filtered by a threshold of the quality score (default: >0.99) reported by the *de novo* software. For each dataset, filtered peptide sequence lists were merged and converted to fasta format using our DeNovo-SIP pipeline (https://git.ufz.de/meb/denovo-sip).

### Identification of isotopically labeled peptides

Mass spectrometric raw files (Supplementary Table S8) were converted into mzML format after peak-picking and encoding in 32-bit precision using msConvert (ProteoWizard v3.0.19046). [85] Next, mzML files were analyzed using the *de novo* peptide databases and for comparison also with the sample-matching protein databases appended with universal contaminants [59] by a customized MetaProSIP workflow [86] based on OpenMS v2.8 [87] on the Galaxy [88] instance of the Helmholtz Centre for Environmental Research – UFZ. In brief, peptides were detected and validated by MS-GF+ [63] using a target-decoy approach, and peptide relative isotope abundances (RIA), as well as labeling ratios (LR), were calculated by MetaProSIP [31]. The precursor mass tolerance was set to ± 10 ppm (for the mock community with spike-ins of ^13^C-labeled *Escherichia coli* [27] and the model of the human gut microbiota [17]) or ± 5 ppm (for the ‘*Candidatus* Kuenenia stuttgartiensis’ enrichment). We allowed a peptide length of 6–100 amino acids, no (for peptide databases) or semi-tryptic (for protein databases) cleavage, PSM-level FDR of 1% (if not stated otherwise), and up to two of the following modifications: variable oxidation of methionine, deamidation of asparagine or glutamine, N-terminal acetylation, carbamylation, or ammonia-loss and fixed C-terminal carbamidomethylation. The weight merge window was set to ± 1.5 (for ^2^H) and ± 5.0 (for ^18^O and ^13^C) in MetaProSIP. Identifications not assigned to a feature during pattern detection and purely natural peptides were included in the report. The peptide-centric output of MetaProSIP was used for further processing. Peptides for which MetaProSIP reported at least one non-natural weight were considered as labeled. The taxonomy and gene ontology (GO) [89] terms of peptides were inferred based on the lowest common ancestor approach by Unipept Desktop v2.0.2 [90] using UniProt 2024.03 [91]. Advanced missed cleavage handling and indistinguishability of leucine and isoleucine were enabled. GO terms of biological processes were mapped to GO slims of the prokaryote subset maintained by the GO Consortium [92] (goslim_prokaryote) using GOATOOLS [93]. Python scripts from our Denovo-SIP pipeline (https://git.ufz.de/meb/denovo-sip) merged results from Unipept and MetaProSIP for downstream analysis.

### Estimation of false discovery proportion

The false discovery proportion (FDP) among peptides identified in at least two samples of a dataset was estimated using an entrapment strategy, as previously described [94]. For this purpose, the original protein sequence databases were augmented with the UniProt reference proteome of the plant *Arabidopsis thaliana* (accession number UP000006548, accession date: 05/14/2025, 27,448 protein sequences), foreign to the investigated samples and frequently used as entrapment [94]. For *de novo* peptide databases, the *A. thaliana* proteome was first *in silico* digested into tryptic peptides with a length of 6–100 amino acids, allowing up to two missed cleavages. A subset of the *A. thaliana* peptides was randomly sampled and appended to the *de novo* peptide databases. The number of sampled entrapment peptides was adjusted to mirror the entrapment-to-target protein ratio (r) of the corresponding reference protein sequence database used in the same dataset. Entrapment-augmented databases were then used for peptide identification via MS-GF+ (see above for parameters). Peptides unambiguously matching entrapment sequences were considered false positives and the false discovery proportion was calculated as previously described [94]:

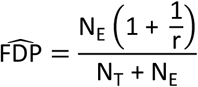

with:

N_E_ = Number of entrapment peptides identified (false positives)

N_T_ = Number of original target peptides identified

*r* = Ratio of entrapment sequences to original target sequences in the database

### Sequence similarity search of unassigned peptides

Labeled peptides that could not be assigned to a lowest common ancestor by Unipept were searched for similar sequences using BLASTP 2.16.0+ [50] with the nr database (accession date: 05/16/2025). Hits with e-value ≥ 0.001 were removed before plotting top-hit species distributions using Blast2GO

6.0.3 [95].

### Taxonomy drop-off rates

The number of taxon-specific peptides decreases as taxonomic resolution increases, resulting in a ‘drop-off’ from higher to lower taxonomic levels. Comparing observed and theoretical drop-off rates within a lineage allows assessing whether the peptide count at a given taxonomic level (e.g., species level) can explain the peptide count at higher taxonomic levels (e.g., phylum level) [51]. By this, drop-off rates can reveal ‘hidden’ organisms not captured by a taxonomic database (here: Unipept). In this manuscript, we use the term ‘superordinate taxa’ to refer to taxonomic ranks above a given taxon. For example, the superordinate taxa of *Escherichia coli* (species) include *Escherichia* (genus), *Enterobacteriaceae* (family), *Enterobacterales* (order), *Gammaproteobacteria* (class), *Pseudomonadota* (phylum), and *Bacteria* (domain).

Labeled peptides were tested for hidden organisms comparing observed with theoretical taxonomy drop-off rates of labeled taxa. To calculate theoretical drop-off rates, we extracted expressed proteins from the proteome of *E. coli* K12 and ‘*Ca.* K. stuttgartiensis’ strain CSTR1 based on previous studies (Table S7 in [60] and Run4_Ecoli268_R3a_0_13C_400ng.pdResult in PXD023693 [96] from [27]). Expressed proteins were *in silico* digested into tryptic peptides with a length of 7–15 amino acids. The detectability of peptides was assessed by DeepMSpeptide [97] and 3,500 detectable peptides from expressed proteins were randomly selected and processed by Unipept as explained above. We accounted for false-positive Unipept annotations by employing a sequence randomization strategy as described previously [51].

### Statistical analysis

Statistical significance was assessed in Python using the `ttest_ind` function from the SciPy library, which performs a two-sample t-test for independent groups. Results were considered significant at p < 0.05 (*), p < 0.01 (**), and p < 0.001 (***).

## Declarations

### Funding

S.K. was funded by the German Research Foundation (DFG) grant number GRK 2032/2. C.W. received funding from the Erasmus+ ICM (KA171) program of the European Commission. Open Access funding was enabled by Project DEAL.

### Availability of data and materials

The raw mass spectrometric proteomics data generated during this study have been deposited to the ProteomeXchange Consortium (http://proteomecentral.proteomexchange.org) via the PRIDE partner repository [96] with the dataset identifier PXD057215 (reviewer token: ejwbd9cUNtVw). Raw mass spectrometry proteomics data reanalyzed during this study is also available via the PRIDE partner repository (Supplementary Table S8). Data obtained from our reanalyses has been deposited to Zenodo (Supplementary Table S8). The Denovo-SIP pipeline’s Python scripts are available via GitLab (https://git.ufz.de/meb/denovo-sip). Any other relevant data will be made available upon request.

### Competing interests

The authors declare that they have no competing interests.

### Authors’ contributions

**Simon Klaes:** Conceptualization, Data curation, Formal analysis, Investigation, Methodology, Project administration, Software, Validation, Visualization, Writing – original draft, Writing – review & editing. **Christian White:** Investigation, Resources, Writing – review & editing. **Lisa Alvarez-Cohen:** Funding acquisition, Resources, Writing – review & editing. **Lorenz Adrian:** Funding acquisition, Resources, Writing – review & editing. **Chang Ding:** Data curation, Formal analysis, Methodology, Project administration, Software, Supervision, Validation, Writing – review & editing

## Acknowledgements

Proteomics raw data published in this study was acquired at UFZ at the Centre for Chemical Microscopy (ProVIS), which is supported by the European Regional Development Fund (EFRE) and the Helmholtz Association. *De novo* sequencing was performed at the High-Performance Computing (HPC) Cluster EVE, a joint effort of both the Helmholtz Centre for Environmental Research – UFZ (http://www.ufz.de/) and the German Centre for Integrative Biodiversity Research (iDiv) Halle-Jena-Leipzig (http://www.idiv-biodiversity.de/). We would like to thank Thomas Schnicke, Ben Langenberg, Guido Schramm, Toni Harzendorf, Tom Strempel, Lisa Schurack, and Christian Krause for maintaining EVE as well as Benjamin Scheer for technical assistance in protein mass spectrometry. We acknowledge the assistance of generative AI (ChatGPT and Claude) in code development and troubleshooting, with all code subsequently reviewed and verified by the authors. Figure 1 was created in BioRender (https://BioRender.com/o64a634).

## Notes

### Competing Interest Statement

The authors have declared no competing interest.

### Summary of Updates

More details on qualitative comparison of de novo peptide databases and (meta)genome derived databases. Method placed more appropriately in the field.

