## Supplementary Information for "*De novo* peptide databases enable protein-based stable isotope probing of microbial communities with up to species-level resolution"

### Supplementary Material

Supplementary Table S1: Sizes of the *de novo* peptide databases employed in this study, with the number of peptides matching the corresponding genome-derived reference protein sequence database indicated in parentheses. The genome-derived reference protein sequence databases were appended with universal contaminants and comprised in total 123,365 protein sequences for dataset PXD024174, 5,344 for PXD057215, and 213,931 for PXD024291.

| **Database size = number of non-redundant *de novo* identified peptides** | | | | | | |
|---|---|---|---|---|---|---|
| **Dataset identifier** | **PXD024174 (Mock community)** | | **PXD057215 (Anammox reactor)** | | **PXD024291 (Human gut model)** | |
| **Score threshold** | Casanovo | PepNet | Casanovo | PepNet | Casanovo | PepNet |
| unfiltered | 290,680 (23,675) | 875,366 (29,429) | 256,559 (7,521) | 303,595 (4,837) | 512,680 (59,600) | 715,031 (43,350) |
| 0.2 | 149,626 (22,743) | 258,054 (23,642) | 82,754 (6,897) | 33,867 (4,773) | 250,323 (55,807) | 225,748 (42,590) |
| 0.4 | 148,847 (22,730) | 175,840 (21,162) | 74,423 (6,896) | 23,321 (4,688) | 247,553 (55,799) | 170,283 (40,904) |
| 0.6 | 139,704 (22,689) | 119,472 (19,003) | 54,101 (6,882) | 16,636 (4,542) | 232,755 (55,705) | 128,653 (39,166) |
| 0.8 | 96,835 (22,426) | 80,093 (17,068) | 29,304 (6,852) | 12,001 (4,370) | 180,762 (55,242) | 97,990 (37,152) |
| 0.85 | 76,673 (22,097) | 71,180 (16,525) | 22,380 (6,781) | 10,960 (4,313) | 153,699 (54,693) | 90,692 (36,575) |
| 0.9 | 53,981 (21,137) | 61,634 (15,894) | 15,785 (6,562) | 9,819 (4,225) | 120,752 (53,113) | 82,718 (35,832) |
| 0.95 | 31,873 (18,813) | 50,450 (15,062) | 9,934 (6,034) | 8,473 (4,100) | 84,176 (48,997) | 72,797 (34,711) |
| 0.99 | 14,700 (13,156) | 35,375 (13,537) | 5,313 (4,644) | 6,605 (3,836) | 48,715 (38,054) | 58,476 (32,424) |

Supplementary Table S2: Number of taxon-specific peptides identified by database search via MS-GF+ using the CS>0.99 database in the PXD024174 dataset.

Supplementary Table S3: Drop-off data from *Escherichia coli*.

| **Labeled peptides** | **Total submissions** | **Root** | **Domain** | **Phylum** | **Class** | **Order** | **Family** | **Genus** | **Species** |
|---|---|---|---|---|---|---|---|---|---|
| in silico | 3,500 | 3,500 | 1,344 | 1,123 | 1,071 | 779 | 659 | 44 | 29 |
| main | 579 | 532 | 190 | 153 | 140 | 85 | 64 | 2 | 2 |
| others | | | 0 | 1 | 7 | 9 | 7 | 9 | 6 |
| random_main | | | 3 | 1 | 0 | 0 | 0 | 0 | 0 |
| random_others | | | 8 | 8 | 9 | 8 | 8 | 6 | 6 |
| main-random_main | | | 187 | 152 | 140 | 85 | 64 | 2 | 2 |
| others-random_others | | | -8 | -7 | -2 | 1 | -1 | 3 | 0 |
| purity [%] | | | 104.5 | 104.8 | 101.4 | 98.8 | 101.6 | 40.0 | 100.0 |
| main-random_main normalised | | | 100.0 | 81.3 | 74.9 | 45.5 | 34.2 | 1.1 | 1.1 |
| *in silico* normalised | | | 100.0 | 83.6 | 79.7 | 58.0 | 49.0 | 3.3 | 2.2 |
| drop off delta [%] | | | | 2.8 | 6.4 | 27.5 | 43.3 | 206.1 | 101.7 |

Supplementary Table S4: Comparison of the number of detected '*Candidatus* Kuenenia stuttgartiensis' strain CSTR1 peptides in unmixed and mixed samples of case study 1.

| **x hydraulic retention time** | | **0** | **1** | | **5.25** | |
|---|---|---|---|---|---|---|
| | | | **unmixed** | **mixed** | **unmixed** | **mixed** |
| '*Candidatus* Kuenenia stuttgartiensis' strain CSTR1 peptides identified by MetaProSIP with genome-derived protein database | | 5,830 | 3,088 | 4,449 | 130 | 3,329 |
| '*Candidatus* Kuenenia stuttgartiensis' strain CSTR1 peptides identified by *de novo* sequencing | Casanovo | 6,695 | 3,628 | 5,120 | 322 | 3,713 |
| | PepNet | 4,336 | 2,035 | 3,171 | 97 | 2,131 |

Supplementary Table S5: Drop-off data from '*Candidatus* Kuenenia stuttgartiensis'.

| Labeled peptides | Total submissions | Root | Domain | Phylum | Class | Order | Family | Genus | Species |
|---|---|---|---|---|---|---|---|---|---|
| in silico | 3,500 | 3,500 | 2,544 | 2,326 | 2,312 | 2,312 | 2,258 | 2,081 | 2,081 |
| main | 1,290 | 1,225 | 894 | 814 | 811 | 811 | 790 | 669 | 669 |
| others | | 0 | 0 | 1 | 1 | 1 | 1 | 1 | 1 |
| random_main | | 339 | 27 | 0 | 0 | 0 | 0 | 0 | 0 |
| random_others | | 0 | 35 | 40 | 40 | 37 | 37 | 35 | 32 |
| main-random_main | | 886 | 867 | 814 | 811 | 811 | 790 | 669 | 669 |
| others-random_others | | 0 | -35 | -39 | -39 | -36 | -36 | -34 | -31 |
| purity [%] | | 100.0 | 104.2 | 105.0 | 105.1 | 104.6 | 104.8 | 105.4 | 104.9 |
| main-random_main normalised | | | 100.0 | 93.9 | 93.5 | 93.5 | 91.1 | 77.2 | 77.2 |
| *in silico* normalised | | | 100.0 | 91.4 | 90.9 | 90.9 | 88.8 | 81.8 | 81.8 |
| drop off delta [%] | | | 0.0 | 2.6 | 2.8 | 2.8 | 2.6 | 6.0 | 6.0 |

Supplementary Table S6. Relative isotope abundance of peptides assigned to gene ontology (GO) terms of biological processes related to subterms of amino acid metabolic processes (GO:0006520), carbohydrate derivative metabolic processes (GO:1901135), and carbohydrate metabolic processes (GO:0005975). P-values were calculated by Student's t-test for the means of two independent samples.

Supplementary Table S7: MS-GF+ runtime comparison between using *de novo* peptide databases (derived from Casanovo after filtering peptides with a quality score threshold > 0.99) and genome-derived protein databases. MS-GF+ was run on the UFZ Galaxy instance using 1 CPU and 57 GB RAM. While genome-derived protein databases were searched with semi-tryptic digestion enabled, *de novo* peptide databases were searched without any digestion enabled. Database size is given in megabytes.

| Dataset identifier | Genome-derived protein database | | CS>0.99 peptide database | |
|---|---|---|---|---|
| | Database size [MB] | Runtime per sample [h] | Database size [MB] | Runtime per sample [h] |
| PXD024174 (Mock community) | 51.8 | 53.5 | 0.4 | 1.0 |
| PXD057215 (Anammox reactor) | 1.6 | 0.9 | 0.3 | 0.3 |
| PXD024291 (Human gut model) | 94.4 | 37.4 | 1.5 | 0.5 |

Supplementary Table S8: Overview of analyzed datasets

| **Dataset** | **Identifier** | **Files analyzed by MetaProSIP workflow** | **Analysis deposited to** |
|---|---|---|---|
| Standard *Escherichia coli* K12 cultures cultivated with 1.07–99% $^{13}C$ [32] | PXD041414 | none | https://zenodo.org/records/15536967?token=eyJhbGciOiJIUzUxMiJ9.eyJpZCI6ImE3YmQzOTE1LTkwMmUtNDUxOS04M2I1LTc3NjQyMTBkNGFjNSIsImRhdGEiOnt9LCJyYW5kb20iOiIyM2NkYjRmMmEyZDAxZWFmZjAxMzcwNzViNDExZGZiZCJ9.P7_Y1CciVFvJLeMB_8oFEVkOAbEMy5W5fE4Lz_9KoKuc8p8-mqYh3Epw2HkK2SwmwuFKwHNcUii7VXFxEZCG9A |
| Mock community with spike-in of $^{13}C$-labeled *Escherichia coli* [27] | PXD024174 | Run1_MockU2_EcoliR1_10_2000ng.raw<br>Run1_MockU2_EcoliR2_10_2000ng.raw<br>Run1_MockU2_EcoliR3_10_2000ng.raw | https://zenodo.org/records/15536771?token=eyJhbGciOiJIUzUxMiJ9.eyJpZCI6ImRlNjRiMTZhLTZlMWEtNDE0MC1hYTlmLTgxZDYwMGFiNjk4OCIsImRhdGEiOnt9LCJyYW5kb20iOiJmZmY0ZmY5YjJlMWFkNTI5YWRjMmE3NDAwNmQ2NmMwNyJ9.xWB5xkIgSib9yDx1nq824XS8lU_UrLuyRHCOgvCNEZDPsUnhaI8wB2JskCwYlBMJ_4mn-TjwfoOwjENud8uJ4g |
| '*Candidatus* Kuenenia stuttgartiensis' strain CSTR1 reactor | PXD057215 | 01_6_27_12C_01.raw<br>02_6_27_12C_02.raw<br>03_6_27_12C_03.raw<br>19_7_01_mix_01.raw<br>20_7_01_mix_02.raw<br>21_7_01_mix_03.raw<br>22_7_18_mix_01.raw<br>23_7_18_mix_02.raw<br>24_7_18_mix_03.raw | https://zenodo.org/records/15536899?token=eyJhbGciOiJIUzUxMiJ9.eyJpZCI6IjdmMGRiMGQ1LWJkMGMtNDBjOC1hOGFiLTZmODM2ZGU0ODE1YyIsImRhdGEiOnt9LCJyYW5kb20iOiI4Y2M3ZGViMWZjZGRiMjRhMWY2NDIzOGYzNzExNWYwNiJ9.NULxsUIzIhoxfapIhgyv-OaY5TPKPsXHpDbQeaEh98Dn84KhFK4nrInCFpLe0x5jZmgMJ9lxzHxYOmTc13YW3w |
| Model of the human distal gut microbiome [17, 27] | PXD024291 | 2017-02-07_RS_Robo_9.raw<br>2017-02-07_RS_Robo_10.raw<br>2017-02-07_RS_Robo_11.raw<br>2017-02-07_RS_Robo_12.raw<br>2017-02-07_RS_Robo_13.raw<br>2017-02-07_RS_Robo_14.raw<br>2017-02-07_RS_Robo_15.raw<br>2017-02-07_RS_Robo_16.raw<br>2017-02-07_RS_Robo_17.raw<br>2017-02-07_RS_Robo_18.raw<br>2017-02-07_RS_Robo_19.raw<br>2017-02-07_RS_Robo_20.raw | https://zenodo.org/records/15537148?token=eyJhbGciOiJIUzUxMiJ9.eyJpZCI6ImZlNWU0MDM4LTRkMjEtNDYxNC1hN2NkLWYzZDY1YjYwMDZiZSIsImRhdGEiOnt9LCJyYW5kb20iOiIyYmY4ZjgwODY5NDY5ZGRkMmI3OTNjZTRlNjdmOTlkYiJ9.CupCCNWTKUZ8ekCcyxHXkmuMNedLw75j0-LJm5ljf3cs24WbrQI38qbFXCy4m7CmadsWMuHIG17_lCvqWzRVxw |

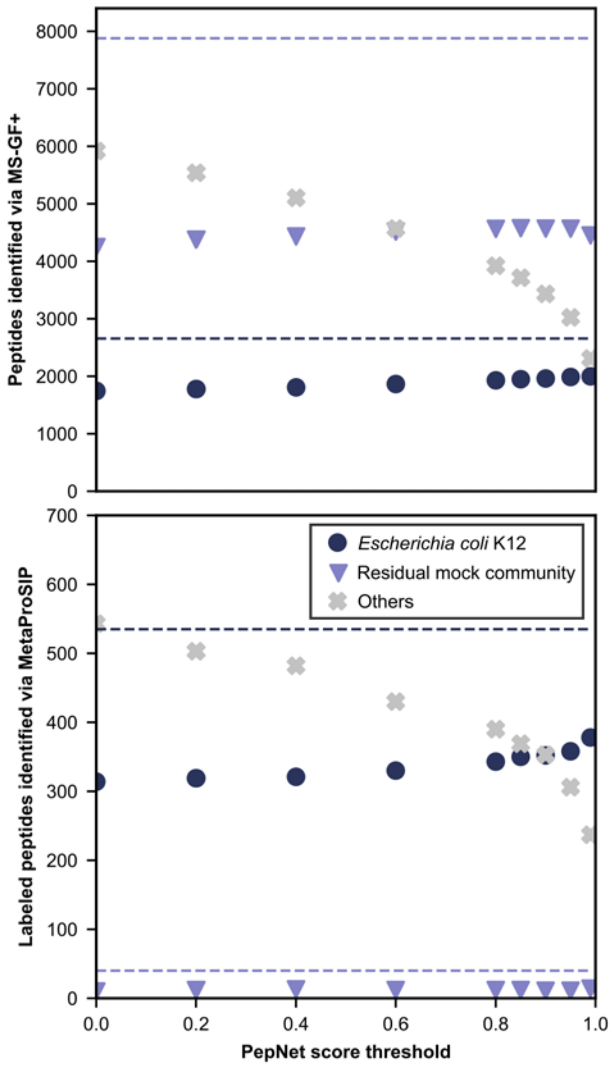


Supplementary Figure S1: Comparison of unlabeled and $^{13}C$-labeled peptide identifications in a mock microbial community spiked with $^{13}C$-labeled *Escherichia coli* K12 cells using MS-GF+ and MetaProSIP with *de novo* peptide databases (1% PSM-level FDR) and a metagenome-derived protein database (2% PSM-level FDR). Raw data was obtained from [27]. The score threshold of >0.99 is highlighted in yellow. Only peptides detected in at least two samples are plotted. **A**: *De novo* peptide databases were generated using Pepnet and filtered at varying quality score thresholds. Identified peptides are color-coded based on their presence in the *E. coli* K12 reference protein database (PRIDE accession number: PXD024285) or common contaminants (navy circles), the metagenome-derived database of the residual mock community (PRIDE accession number: PXD006118, purple triangles), or neither (grey crosses). Dashed lines represent identifications from the metagenome-derived database.

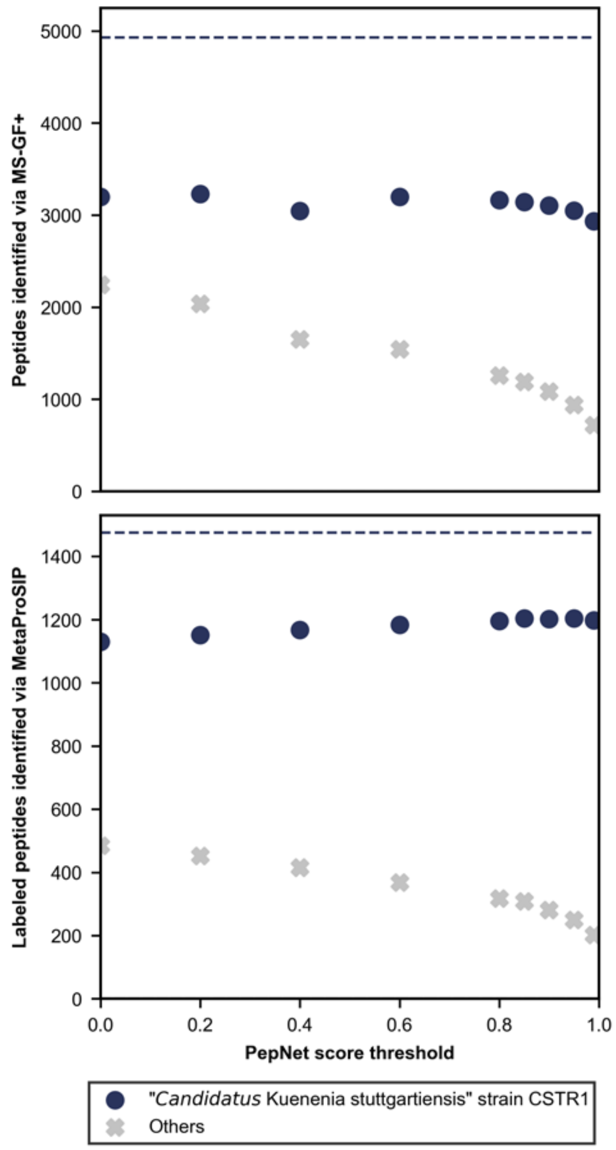


Supplementary Figure S2: Comparison of unlabeled and $^{13}C$-labeled peptide identifications using MS-GF+ and MetaProSIP in an enrichment of '*Candidatus* Kuenenia stuttgartiensis' strain CSTR1 from a continuous flow reactor fed with $^{13}C$-bicarbonate with *de novo* peptide databases (1% PSM-level FDR) and a genome-derived protein database (2% PSM-level FDR). The score threshold of >0.99 is highlighted in yellow. Only peptides detected in at least two samples are plotted. **A**: *De novo* peptide databases were generated using PepNet and filtered at varying quality score thresholds. Navy circles represent peptide identifications matching the genome-derived database (NCBI: CP049055.1) or universal contaminants; grey crosses represent unmatched peptides. Dashed lines show identification counts using the genome-derived database.

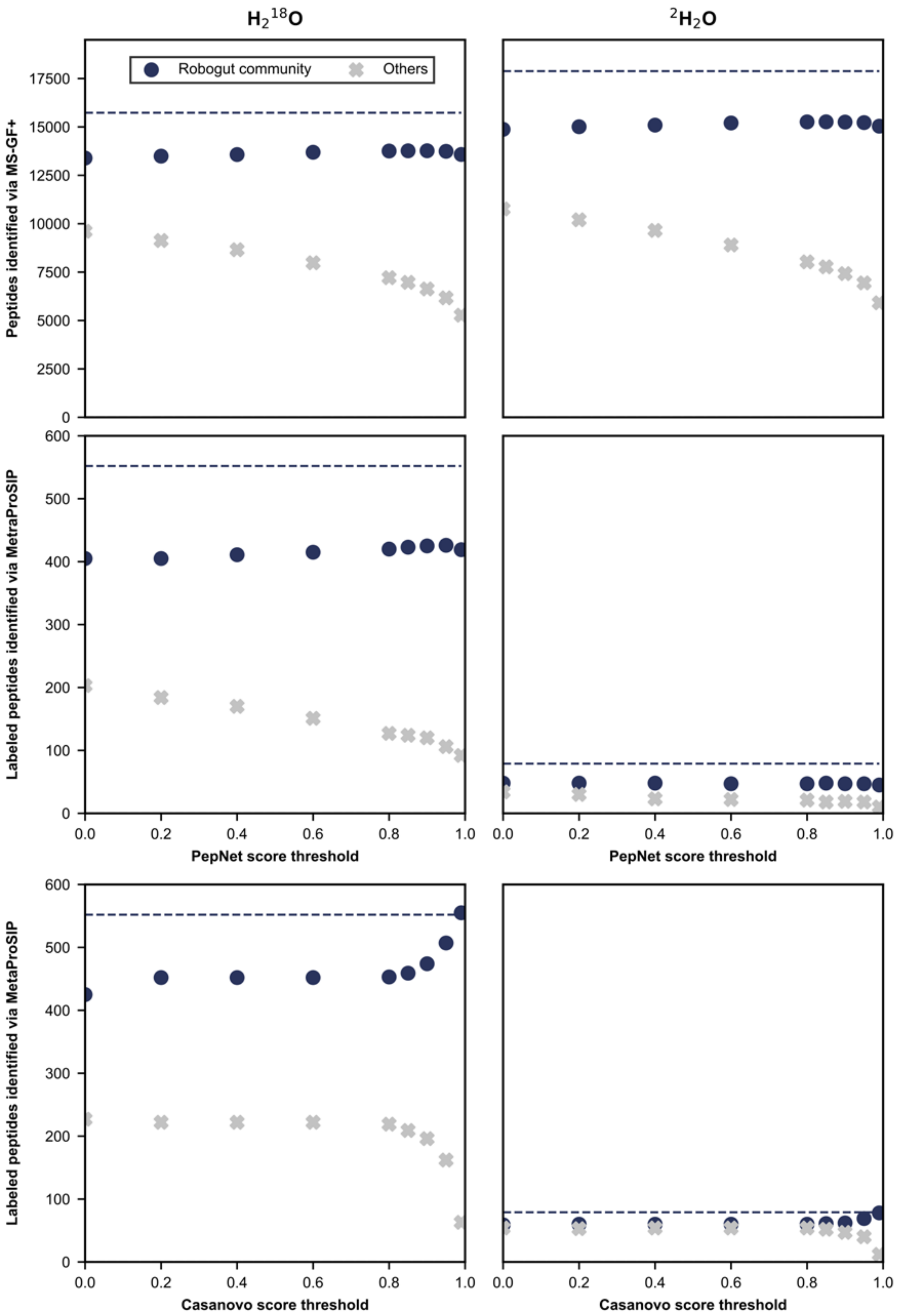

$H_2{}^{18}O$
$^2H_2O$
Robogut community
Others
Peptides identified via MS-GF+
Labeled peptides identified via MetraProSIP
PepNet score threshold
Casanovo score threshold

Supplementary Figure S3: Comparing *de novo* peptide databases for identifying unlabeled and $^{2}H/^{18}O$-labeled peptides via MS-GF+ and MetaProSIP (1% PSM-level FDR) in a model of the human gut microbiome (Robogut community) cultivated with $^{2}H_2O$ or $H_2{}^{18}O$, respectively, and obtained from [17]. *De novo* peptide databases were assembled using PepNet or Casanovo and filtering identifications with different thresholds for the quality score. Navy circles denote the number of identified peptides present in the genome-derived reference database or universal contaminants. Grey crosses denote the number of identified peptides absent in the genome-derived reference database and universal contaminants. Dashed lines represent the number of peptides identified using the genome-derived reference database appended with universal contaminants. Only peptides detected in at least two samples are plotted.

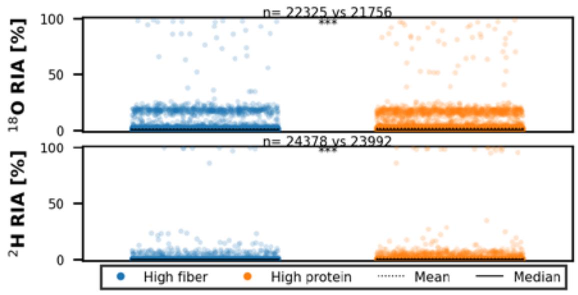


Supplementary Figure S4: Distribution of the $^{18}O$ and $^{2}H$ relative isotope abundance (RIA) in detected peptides identified in a model of the human gut microbiome cultivated in either a high fiber or high protein medium in the presence of either $H_2{}^{18}O$ or $^{2}H_2O$. RIA values are shown for peptides detected in at least two replicates of both media. 'n' denotes the number of peptides identified across triplicates. Statistically significant differences in RIA between media are indicated with '***' based on Student's t-test for the means of two independent samples with $p < 0.001$.

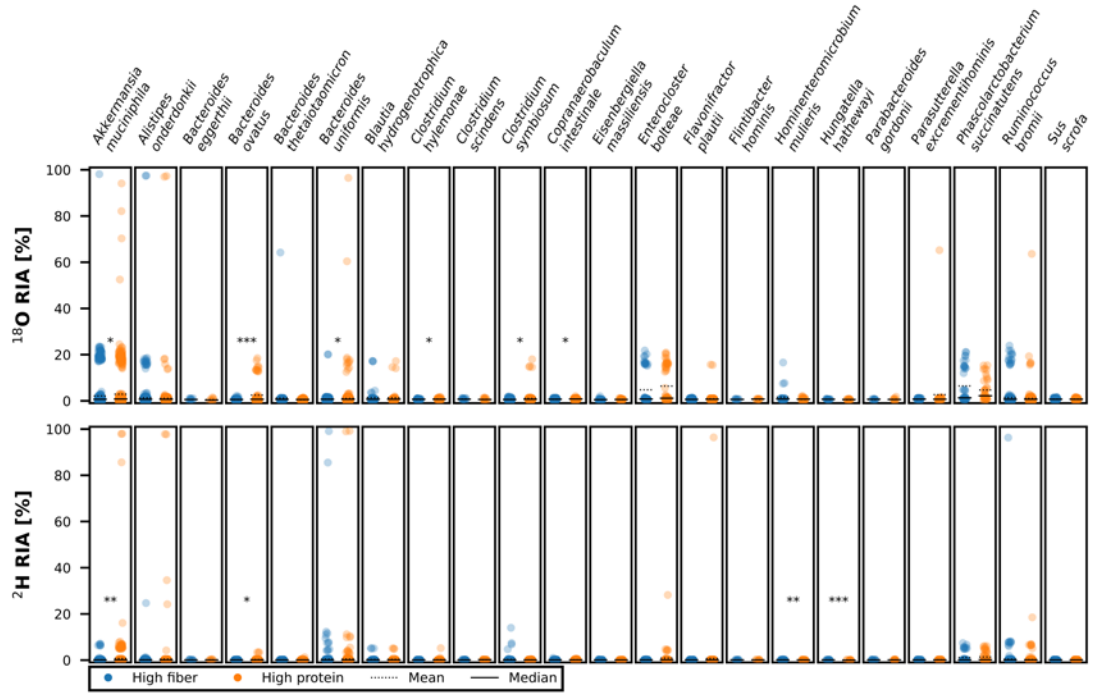


Supplementary Figure S5: Figure 8 without removal of outliers.

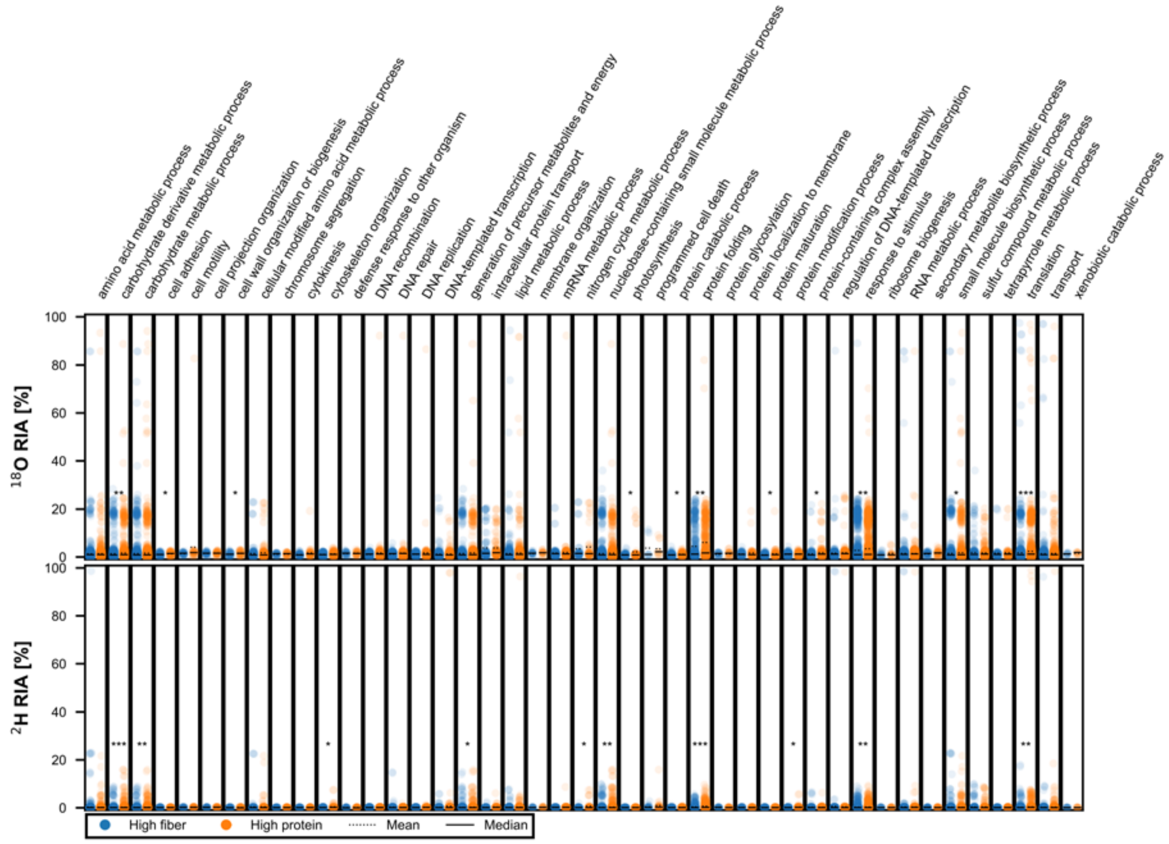


Supplementary Figure S6: Figure 9 without removal of outliers.
